# Sociosexual behavior requires both activating and repressive roles of Tfap2e/AP- 2ε in vomeronasal sensory neurons

**DOI:** 10.1101/2022.01.14.476379

**Authors:** Jennifer M. Lin, Tyler A. Mitchell, Megan Rothstein, Alison Pehl, Ed Zandro M. Taroc, Raghu Ram Katreddi, Katherine E. Parra, Damian G. Zuloaga, Marcos Simoes-Costa, Paolo E. Forni

## Abstract

Neuronal identity dictates the position in an epithelium, and the ability to detect, process, and transmit specific signals to specified targets. Transcription factors (TFs) determine cellular identity via direct modulation of genetic transcription and recruiting chromatin modifiers. However, our understanding of the mechanisms that define neuronal identity and their magnitude remains a critical barrier to elucidate the etiology of congenital and neurodegenerative disorders. The rodent vomeronasal organ provides a unique system to examine in detail the molecular mechanisms underlying the differentiation and maturation of chemosensory neurons. Here we demonstrated that the identity of postmitotic/maturing vomeronasal sensory neurons (VSNs), and vomeronasal dependent behaviors can be reprogrammed through the rescue of tfap2e/AP-2ε expression in the AP-2ε mice, and partially reprogrammed by inducing ectopic AP-2ε expression in mature apical VSNs. We suggest that the transcription factor AP-2ε can reprogram VSNs bypassing cellular plasticity restrictions, and that it directly controls the expression of batteries of vomeronasal genes.

**GRAPHICAL ABSTRACT:** 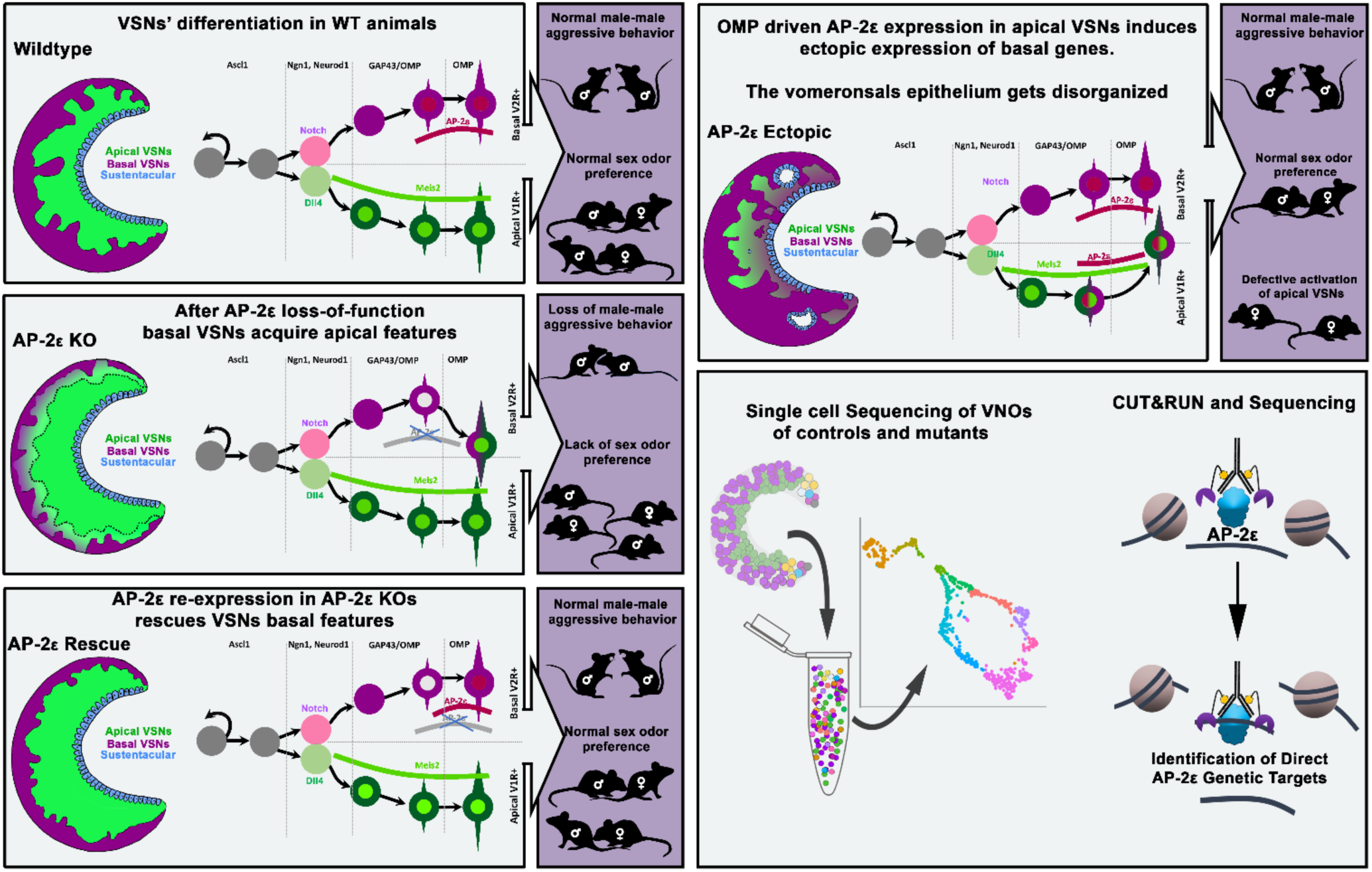

## INTRODUCTION

Neuronal differentiation is controlled by the selective expression of transcription factors (TFs), chromatin modifiers, and other regulatory factors that reduce cellular plasticity. During neuronal differentiation terminal selectors can activate identity-specific genes that define functional properties specific to a particular neuronal type. However, reprograming postmitotic neurons by ectopically expressing terminal selectors in *C. elegans* suggests that the “reprogrammability” of neurons is progressively lost during postembryonic life (Patel and Hobert, 2017; Patel et al., 2012; Rahe and Hobert, 2019). This reduction in cellular plasticity may arise from chromatin modifications that prevent the activation of alternative differentiation programs. However, sensory neurons of the vomeronasal organ in rodents can undergo some postnatal reprogramming following the aberrant expression of transcription factors (Lin et al., 2018).

The accessory olfactory system (AOS) contains the vomeronasal organ (VNO), which is primarily responsible for detecting odors and chemosignals that trigger social and sexual behaviors (Trouillet et al., 2019; Trouillet et al., 2021). The vomeronasal sensory epithelium of rodents is mainly composed of vomeronasal sensory neurons (VSNs). The VSN populations selectively express only one or two receptors encoded by the two vomeronasal receptor (VR) gene families: V1R and V2R (Dulac and Axel, 1995; Herrada and Dulac, 1997; Matsunami and Buck, 1997; Ryba and Tirindelli, 1997). V1R and V2R-expressing neuronal populations each detect distinct chemosignals, induce different innate behaviors, show distinct localization patterns in the VNO, and project to specific areas of the accessory olfactory bulb (AOB) (Cloutier et al., 2002; Dulac and Axel, 1995; Isogai et al., 2011; Katreddi and Forni, 2021; Mohrhardt et al., 2018; Mombaerts et al., 1996; Stowers et al., 2002). The V2R-expressing neurons localize to the basal portions of the vomeronasal epithelium (VNE) and around the vasculature (Naik et al., 2020), while V1R-expressing neurons localize to the apical part. Basal and apical VSNs continually regenerate from common pools of Achaete Scute like-1 (Ascl-1)-positive neural progenitor cells (NPCs) localized in the lateral and basal margins of the VNE (Cau et al., 1997; de la Rosa-Prieto et al., 2010; Katreddi and Forni, 2021; Martinez-Marcos et al., 2000; Murray et al., 2003). However, we are only starting to understand how the apical and basal VSN cell differentiation programs are initiated and which factors aide in maintaining apical and basal neuronal identity (Enomoto et al., 2011; Katreddi et al., 2022; Lin et al., 2018; Naik et al., 2020; Oboti et al., 2015).

Establishing functional basal and apical VSNs is crucial for intra- and interspecies social interactions in rodents. Deficits in basal neuron functionality prevented sex discrimination, reduced male-male and maternal aggressive behaviors, and inhibited the detection of predator odors (Chamero et al., 2011; Stowers et al., 2002).

Transcription factors can drive cellular processes that control the expression of genes defining their cellular and functional identity. The AP-2 family of transcription factors is comprised of 5 members AP-2α, AP-2β, AP-2γ, AP-2δ and AP-2ε, which are encoded by distinct genes (TFAP2A, TFAP2B, TFAP2C, TFAP2D, TFAP2E) (Eckert et al., 2005; Pellikainen and Kosma, 2007; Wankhade et al., 2000). AP-2 family members play critical roles during development, such as contributing to neural crest differentiation (Luo et al., 2020; Rothstein and Simoes-Costa, 2020), cell specification, limb development, and organogenesis (Bassett et al., 2012; Chambers et al., 2019; Kantarci et al., 2015). Some AP-2 family members may have pioneer factor properties (Fernandez Garcia et al., 2019; Rothstein and Simoes-Costa, 2020; Seberg et al., 2017; Williams et al., 2009).

Aside from AP-2ε, Notch signaling and Bcl11b control Gαo+ VSNs’ differentiation, homeostasis, and survival (Enomoto et al., 2011; Katreddi et al., 2022). We previously proposed that AP-2ε, which is only expressed after the apical and basal VSN dichotomy is established, is necessary for further specification of basal VSN identity (Lin et al., 2018). Using mice expressing non-functional AP-2ε, we discovered that VSNs can still acquire the Gαo+/basal identity; however, these VSN have reduced survival and can acquire some Gαi2+/apical VSNs molecular features over time (Lin et al., 2018). While we examined the role of AP-2ε in maintaining cellular identity and homeostasis of the vomeronasal epithelium, critical outstanding questions remain unresolved. What role does this transcription factor actively play to control the basal genetic program? What is the extent of cellular plasticity in differentiated neurons in mammals (Patel and Hobert, 2017; Rahe and Hobert, 2019)?

Here, we aimed to understand 1) if AP-2ε functions as a terminal selector factor for basal VSNs, 2) how much cellular plasticity postmitotic neurons retain once differentiated, and 3) to what extent genetic dysregulation in mature VSNs translates into behavioral changes. We generated a Cre inducible mouse line, where we inserted the mTfap2e/AP-2ε gene into the ROSA26 locus. Using this knock-in mouse line, we could 1) rescue the AP-2ε KO’s VNO morphology and functionality and 2) ectopically express AP-2ε in maturing Gαi2+/apical VSNs. This approached enable us to assess its ability to reprogram differentiated apical VSNs to basal VSNs. By combining histological analyses, behavioral assessments, and single-cell RNA sequencing (scRNA-seq) analysis, we examined whether AP-2ε functions as a master regulator to reprogram differentiated neurons and alter animal behaviors. In addition, we used CUT&RUN (Skene et al., 2018) to identify direct genetic targets of AP-2ε that controls the basal VSNs identity program. Overall, we suggest that AP-2ε partially functions as a terminal selector by activating some basally enriched genes while simultaneously suppressing specific apically enriched genes.

## RESULTS

### Transcriptome differences between apical and basal VSNs

Using single cell RNA-sequencing (scRNA-Seq) on VNOs from OMPCre+ control mice at P10, we identified key features of VSNs based on the expression plots. We then clustered single cells into representative Uniform Manifold Approximation and Projections (UMAPs) (Figure 1). Achaete-Scute Family BHLH Transcription Factor 1 (Ascl1), (Figure 1A), Neurogenin1 (Neurog1) (Figure 1B), and NeuroD1 (Figure 1C) expression identified proliferative VSN progenitors (Katreddi and Forni, 2021). We determined that the dichotomy of apical-basal differentiation begins when the cells transition from Neurog1 to NeuroD1 expression, and during the NeuroD1 phase (Figure 1B,C)(Katreddi et al., 2022).

**Figure 1:**
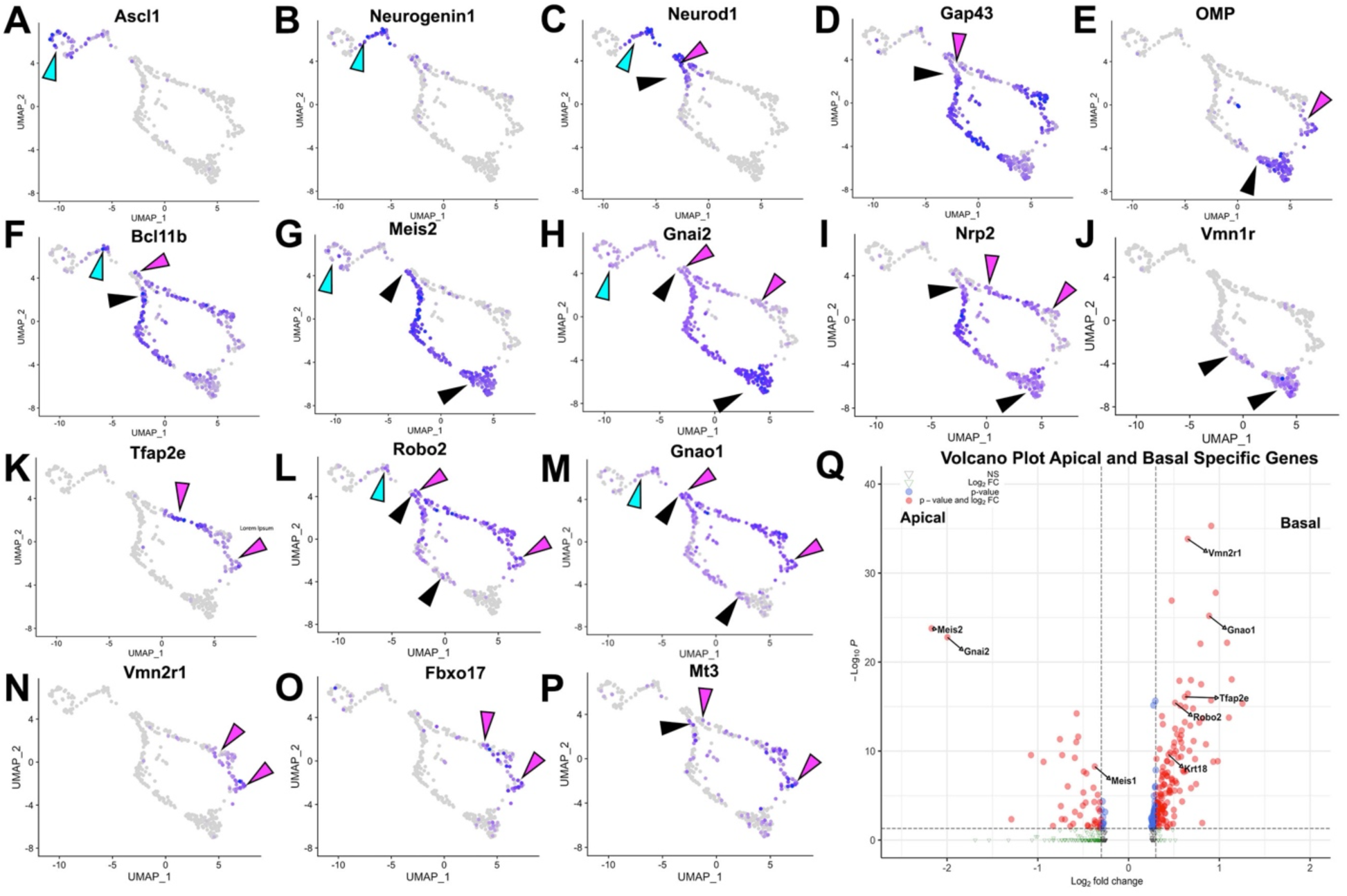
Analysis of single cell sequencing data of the vomeronasal organ. P10 male controls shows the developing vomeronasal neurons as they progress through A-C) neurogenesis (Ascl1, Neurog1, Neurod1), D-F) maturation (Gap43, OMP, Bcl11b), G-J) apical VSNs’ differentiation/maturation (Meis2, Gnai2, Nrp2, Vmn1r) and K-P) basal VSNs’ differentiation/maturation (Tfap2e, Robo2, Gnao1, Vmn2r1, Fbxo17, Mt3). A) Ascl1 is expressed by transiently amplifying progenitor cells (cyan arrow), which transition into the immediate neuronal precursors that turn on pro-neural genes (cyan arrow) Neurog1 (B) and Neurod1 (C) and turn off as the precursors turn into immature neurons (black and magenta). D) Immature neurons express Gap43, which persists in both apical (black arrow) and basal (magenta arrow) branches, until it declines as neurons begin to reach maturity and express E) OMP in mature apical (black arrow) and mature basal (magenta arrow) VSNs. Gap43 and OMP expression briefly overlap, as the neurons transition to a fully differentiated mature stage. F) Bcl11b mRNA expression is found in both apical and basal VSNs but at different developmental timepoints. Bcl11b is found in committed basal precursors near the establishment of the apical/basal dichotomy (magenta arrow) but is not found until later in apical VSN development (black arrow) G) Meis2 mRNA expression is found in the apical branch (black arrows) and even in early neurogenesis stages (cyan arrow), and their expression does not overlap. H) Gnai2 expression starts in immature apical and basal VSNs. However, its expression increases in mature apical VSNs but fades in mature basal VSNs. I) Nrp2 expression is only retained in mature apical VSNs. J) Vmn1r is only expressed in maturing/mature apical VSNs. K) AP-2ε is expressed by the basal branch (magenta arrows). L) Robo2 expression is only retained in mature basal VSNs. M) Gnao1 expression starts in immature apical and basal VSNs. However, its expression increases in mature basal VSNs but fades in mature apical VSNs. N) Vmn2r1 is restricted to maturing/mature basal VSNs. O) Fbxo17 is mainly expressed in mature/maturing basal VSNs. P) Mt3 is mainly expressed in mature/maturing basal VSNs. Q) Enhanced volcano plot. Differential gene expression between apical and basal branches of VSNs. Apical specific genes (55 genes) trend left and basal specific genes (187 genes) trend right. Significance defined as Log_2_-Fold Change > 0.3 and Adjusted p-value ≤ 0.05.

The later stages of apical and basal VSN maturation were marked by the expression of Gap43 (Figure 1D) in immature VSNs and OMP (Figure 1E) in more mature VSNs (Katreddi and Forni, 2021).

Consistent with prior reports, the mRNA for the transcription factor Bcl11b (Figure 1F) was found to be expressed in Neurog1+/NeuroD1+ (Figure 1B,C) precursors and in differentiating apical and basal VSNs. While Bcl11b was expressed as a continuum along the basal differentiation trajectory, in the apical neurons, Bcl11b was not expressed until later stages of maturation (Figure 1F) (Enomoto et al., 2011; Katreddi et al., 2022). The transcription factor Meis2 was expressed in progenitor cells as well as in apical VSNs’ differentiating neurons (Figure 1G). These also express Gnai2/ Gαi2, Nrp2 and V1rs at more mature stages (Figure 1I-K). On the other side, mRNA for the transcription factors Tfap2e/AP-2ε (Figure 1K) was found to be expressed, in line with our prior works (Katreddi et al., 2022; Lin et al., 2018), in differentiating/maturing and mature basal VSNs. The basal VSNs’ differentiation trajectory was further confirmed by the expression of known V2r/basal VSNs’ markers such as Robo2, Gnao1/Gαo, and Vnm2r receptors (Figure 1K-N). In addition, our transcriptome analysis revealed significant enrichments (q<0.05, Figure 1Q, Supplementary Table S1) of several previously unreported genes in either apical or basal VSNs. Fbxo17, Mt3 (Figures 1O,P; S5) and Keratin18 (Krt18), were among the genes that we found enriched in maturing basal VSNs.

### Inducible R26AP2ε rescues basal VSNs in AP-2ε KOs

We hypothesized that Tfap2e (AP-2ε) can control gene expression during basal VSN maturation. So, we generated a new Cre inducible mouse line (B6.Cg-Gt(ROSA)26Sor^tm(CAG-mTfap2e)For^). We inserted a lox-P-flanked stop cassette to prevent the transcription of a CAG promoter driven murine Tfap2e gene, which was knocked into the first intron of the Gt(ROSA)26Sor locus (Figure 2A). We refer to this as R26AP2ε. AP-2ε expression is normally restricted to basal regions of the VNO with higher expression levels of AP-2ε in the neurogenic marginal zones (Enomoto et al., 2011; Lin et al., 2018; Naik et al., 2020)(Figures 1; 2B,B’). To test our Cre inducible AP-2ε line, we performed anti-AP-2ε immunostaining on wild-type controls, AP-2ε^Cre/Cre^ (AP-2ε^Null^)(Feng et al., 2009; Lin et al., 2018), and AP-2ε^Cre/Cre^/R26AP2ε (AP-2ε^Rescue^) mice (Figure 2B-D). As expected, (Lin et al., 2018), wild-type mice showed AP-2ε immunoreactivity in the basal regions of the VNE with strong immunoreactivity in the neurogenic regions (Figure 2B, B’). However, in AP-2ε^Null^ mice, where Cre was knocked into the DNA binding domain of AP-2ε (Feng et al., 2009; Feng and Williams, 2003), we observed faint AP-2ε cytoplasmic immunoreactivity limited to the most marginal zones of the VNO and no immunoreactivity in the rest of the neuroepithelium (Figure 2C,C’). AP-2ε^Rescue^ mice showed restored AP-2ε immunoreactivity in the basal region of the VNE (Figure 2D). However, we observed that the AP-2ε expression pattern and immunoreactivity were not identical to controls in the neurogenic regions. In AP-2ε^Rescue^ mice, we observed no AP-2ε immunoreactivity at the tips of the neurogenic niche in the VNE (Figure 2D’), suggesting a delayed AP-2ε expression after AP-2εCre mediated recombination compared to controls (Figure 2B’,D’).

**Figure 2:**
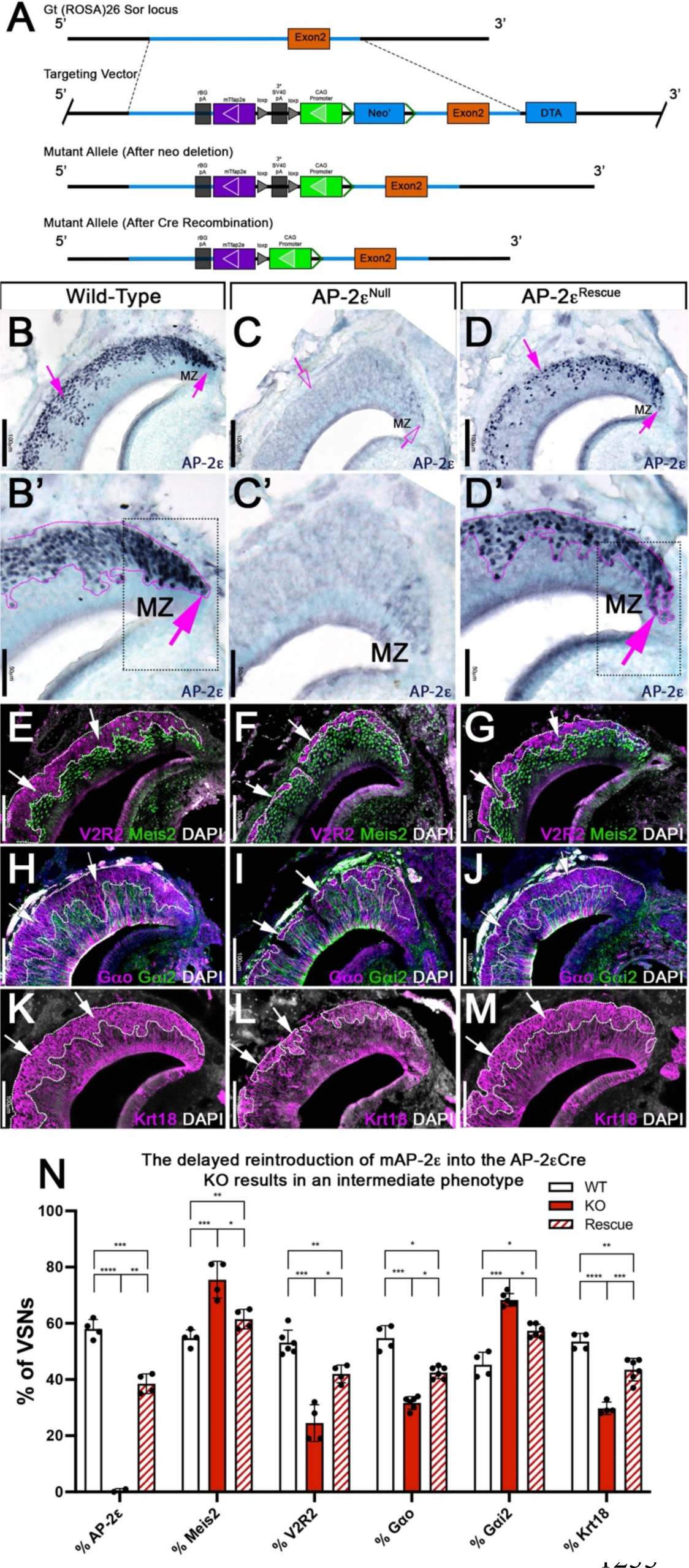
R26AP2ε mouse line mouse generation and characterization of rescued AP-2ε expression in AP-2ε^Null^ mice. A) Knock-in strategy through homologous recombination to generate the R26AP2ε mouse line. The CAG-loxP-stop-loxP-mouse-Tfap2e cassette as integrated into the first intron of Rosa26. B-D’) Immunohistochemistry on P21 wild-type (B, B’), AP-2ε^Null^ (C,C’), and AP-2ε^Rescue^ (D,D’) mice against AP-2ε. B,B’) In WT mice AP-2ε expression is in the marginal zones (MZ) and in the basal regions of the VSNs. C,C’) In AP-2ε^Null^ mice, some AP-2ε immunoreactivity is observed in the MZ but lost in the central regions where more mature neurons reside. D,D’) In the AP-2ε^Rescue^, AP-2ε is expressed in the basal region, but with less intensity and density at the MZ and in central regions compared to WT controls. E-G) Immunostainings against V2R2 (magenta) and Meis2 (green) counterstained with DAPI (white). H-J) Immunostainings against Gαo (magenta) and Gαi2 (green) counterstained with DAPI). K-M) Immunostainings against Krt18 (magenta) counterstained with DAPI (white). N) Quantifications of the percentage of VSNs expressing apical/basal markers in WT, AP-2ε^Null^ and AP-2ε^Rescue^ mice. AP-2ε^Null^ mice show a dramatic reduction in basal VSNs and apical VSNs occupy most of the epithelium. The AP-2ε^Rescue^ has an intermediate phenotype between WT and AP-2ε^Null^ mice, where the VNE contains more basal VSNs than in the AP-2ε^Null^ mice but does not reach the equivalency of the WT (p<0.05=*, p<0.01=**, p<0.001=***, p<0.0001=****). N=4 for WT in % AP-2ε, % meis2, % Gαo, % Gαi2, and % Krt18. N=6 for WT in % V2R2. N=2 for AP-2ε^Null^ in % AP-2ε. N=4 for AP-2ε^Null^ in % Meis2, % V2R2 and % Krt18. N=6 for AP-2ε^Null^ in % Gαo and % Gαi2. N=4 for AP-2ε^Rescue^ in % AP-2ε, % Meis2 and % V2R2. N=6 for AP-2ε^Rescue^ in % Gαo, % Gαi2, and % Krt18.

By analyzing the expression of the basal markers downregulated in AP-2ε KOs (Lin et al., 2018), such as V2R2 (Figure 2E-G) and Gαo (Figure 2H-J), we confirmed restored expression in the rescued KOs. Cell quantifications indicated a significant increase in the number of basal cells expressing basal markers in AP-2ε^Rescue^ compared to AP-2ε^Null^ mice, though the number of basal VSNs in the AP-2ε^Rescue^ was smaller when compared to controls (Figure 2N). Keratin18 (Krt18) is normally enriched in basal neurons (Figure 1Q). Immunohistochemistry confirmed Krt18 protein expression in the basal territories of the VNO (Figure 2K). We observed reduced Krt18 immunoreactivity in AP-2ε^Null^ mice and restored expression in AP-2ε^Rescue^ mice (Figure 2L-N). However, in AP-2ε^Rescue^ mice, Krt18 still showed lower expression levels than in WT mice (Figure 2N). Taken together, we conclude that exogenous AP-2ε in postmitotic VSNs can partially rescue the expression of basal VSN markers in AP-2ε^Nulls^ mice. Rescue of the AP-2ε^null^ phenotype indicates that our inducible R26AP2ε mouse line is a suitable model for conditional expression of functional AP-2ε.

### Re-expressing AP-2ε in AP-2ε mice rescues social behaviors

The specification and organization of VSNs and their respective circuit assembly in the AOB are essential to trigger a variety of social and sexual behaviors (Chamero et al., 2011; Chamero et al., 2007; Stowers et al., 2002; Trouillet et al., 2019) . We speculated that AP-2ε^Null^ mice could not discriminate between urine of different sexes. Thus, we performed an odorant preference test. In this test, individual mice were simultaneously presented with male and female whole urine for a two-minute period (Figure 3A). Wild-type male mice showed a significant preference for urine from the opposite sex (Figure 3B)(Pankevich et al., 2004; Stowers et al., 2002). However, AP-2ε^Null^ male mice did not display significant preference for female urine, confirming a loss of function (LOF) of basal VSNs (Lin et al., 2018) and consequently a reduced ability to discriminate between urine from either sex (Figure 3B)(Pankevich et al., 2004; Stowers et al., 2002). However, AP-2ε^Rescue^ male mice showed a significant preference for female urine similar to WT controls (Figure 3B).

**Figure 3:**
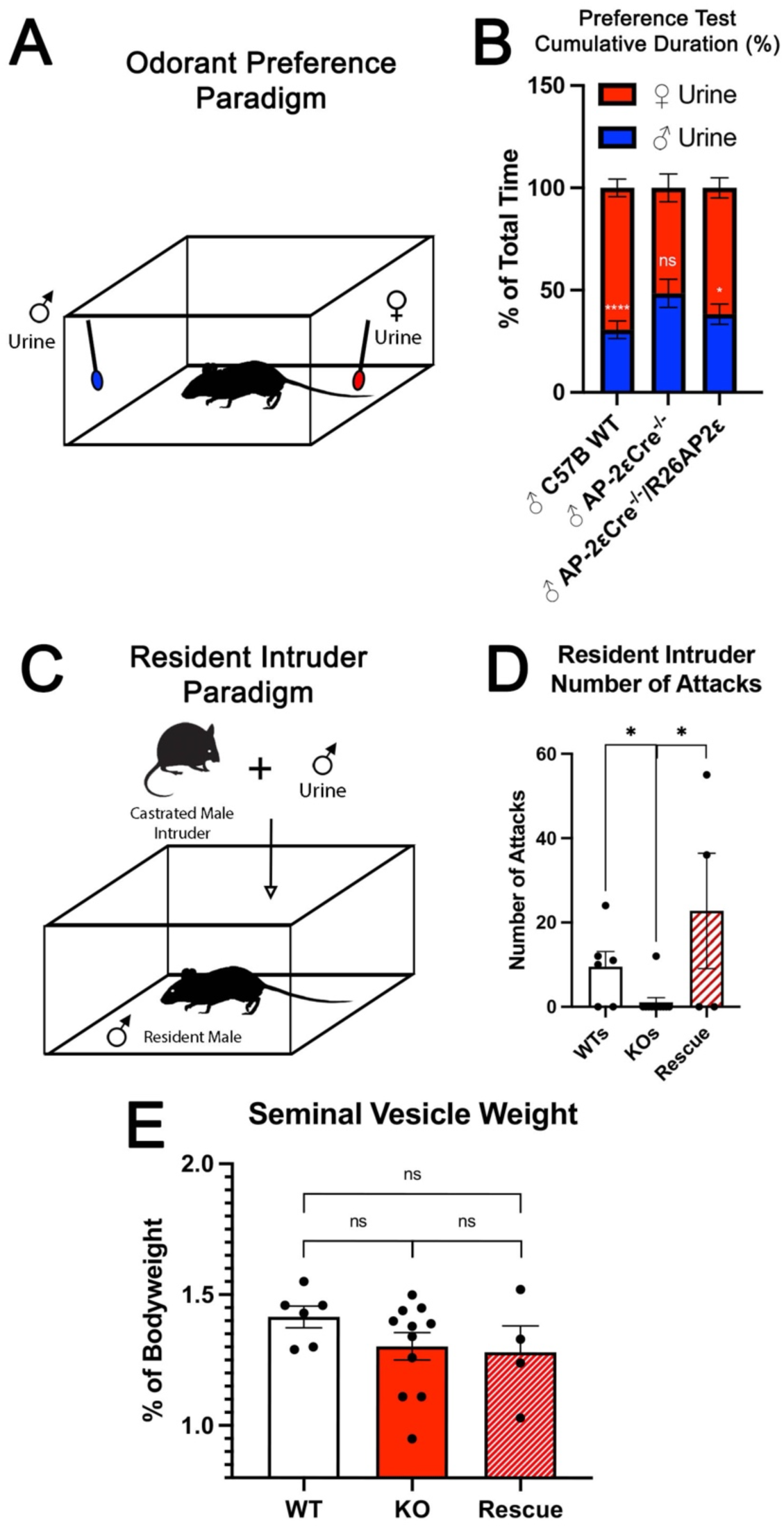
Territorial aggression and sex preference depends on AP-2ε expression in mice. A) The odorant preference paradigm where cotton swabs with either male or female whole urine are placed on opposite ends of a test cage and the amount of time spent smelling each odorant is measured. B) Male WT mice spent significantly more time investigating female odorants than male odorants. Preference for female odorants is lost in AP-2ε KO male mice but restored in AP-2ε rescue mice. (p<0.05=*, p<0.01=**, p<0.001=***). N=6 for WT, n=11 for AP-2ε^Null^ and n=4 for AP-2ε^Rescue^. C) Male-male aggression was evaluated using the Resident intruder paradigm D) WT mice display aggressive behaviors toward male intruders and number of attacks were quantified. AP-2ε Null mice attacked intruders significantly less than WT mice. However, AP-2ε Rescue mice showed significantly more aggressive behaviors that is not significantly different than the WT male mice. (p<0.05=*). N=6 for WT, n=11 for KO, n=4 for rescue. E) Seminal vesicle weight was not significantly different across all genotypes when normalized to bodyweight. N=6 for WT, n=11 for KO, n=4 for rescue.

To further investigate the behavioral outcome of AP-2ε LOF and AP-2ε re-expression in AP-2ε^Null^ mice, we performed a resident intruder assay for intermale aggression (Figure 3C)(Chamero et al., 2011; Montani et al., 2013; Stowers et al., 2002). Wild-type male mice showed aggressive behaviors toward intruders upon detecting male specific odorants (Figure 3D). Most AP-2ε^Null^ mice did not attack the intruder (Figure 3D). Yet, when male AP2ε^Cre/Cre^/R26AP2ε^+/-^ (AP-2ε^Rescue^) mice were exposed to male intruders, they displayed aggressive behavior similar to that of controls (Figure 3D). We measured the mass of the seminal vesicles from each genotype to rule out any changes in general androgen levels, which may explain any potential behavioral differences (Zuloaga et al., 2007). We found no significant differences when the seminal vesicle weights were normalized to the total body weight of each mouse (Figure 3E). Taken together, these data suggest re-expression of AP-2ε in KO mice can reestablish the functional properties of basal VSNs.

### Ectopic expression of AP-2ε in mature apical VSNs increases the expression of basal enriched genes

Several AP-2 family members have been proposed to have pioneer activity (Fernandez Garcia et al., 2019; Rothstein and Simoes-Costa, 2020; Seberg et al., 2017; Williams et al., 2009). We tested whether ectopic AP-2ε expression can alter the transcriptomic profile of maturing neurons. Olfactory marker protein (OMP) is an accepted marker for postmitotic/maturing olfactory and vomeronasal sensory neurons (Buiakova et al., 1994; Enomoto et al., 2011; Farbman and Margolis, 1980). By analyzing our scSeq data from OMPCre^+/-^ mice, we confirmed that OMP mRNA expression can be detected in maturing apical neurons shortly after the apical basal dichotomy is established (Figure 1E). Thus, we used an OMPCre mouse line to drive expression of AP-2ε (OMPCre^+/-^ /R26AP2ε^+/-^) in all olfactory and vomeronasal neurons. In this manuscript we will often refer to OMPCre^+/-^/R26AP2ε^+/-^ as ectopic mutants. Notably, both apical and basal VSNs express several known Tfap2 cofactors, including Cited2 and p300/CBP (Bamforth et al., 2001; Bragança et al., 2003; Eckert et al., 2005) suggesting that both VSN populations are molecularly competent for functional AP-2ε transcriptional activity (Figure S1).

Immunostaining against AP-2ε showed no immunoreactivity in the main olfactory epithelium (OE) of control animals (Figure S2A). However, in OMPCre^+/-^/R26AP2ε^+/-^ mutants, the OE expressed immunodetectable AP-2ε (Figure S2C). OMPCre^+/-^ controls and OMPCre^+/-^/R26AP-2ε^+/-^ mutants displayed comparable gross morphology of the OE with no ectopic V2R immunoreactivity (Figure S2B,D).

In the VNO of OMPCre^+/-^ controls, AP-2ε was found only in cells in the basal territory (Figure 4A, A1). However, in OMPCre^+/-^/R26A-P2ε^+/-^ mice, we found that virtually all the VSNs expressed AP-2ε (Figure 4D, D1). In these mutants we also observed AP-2ε expression in sparse sustentacular cells lining the lumen of the VNO (Figure 4D).

**Figure 4:**
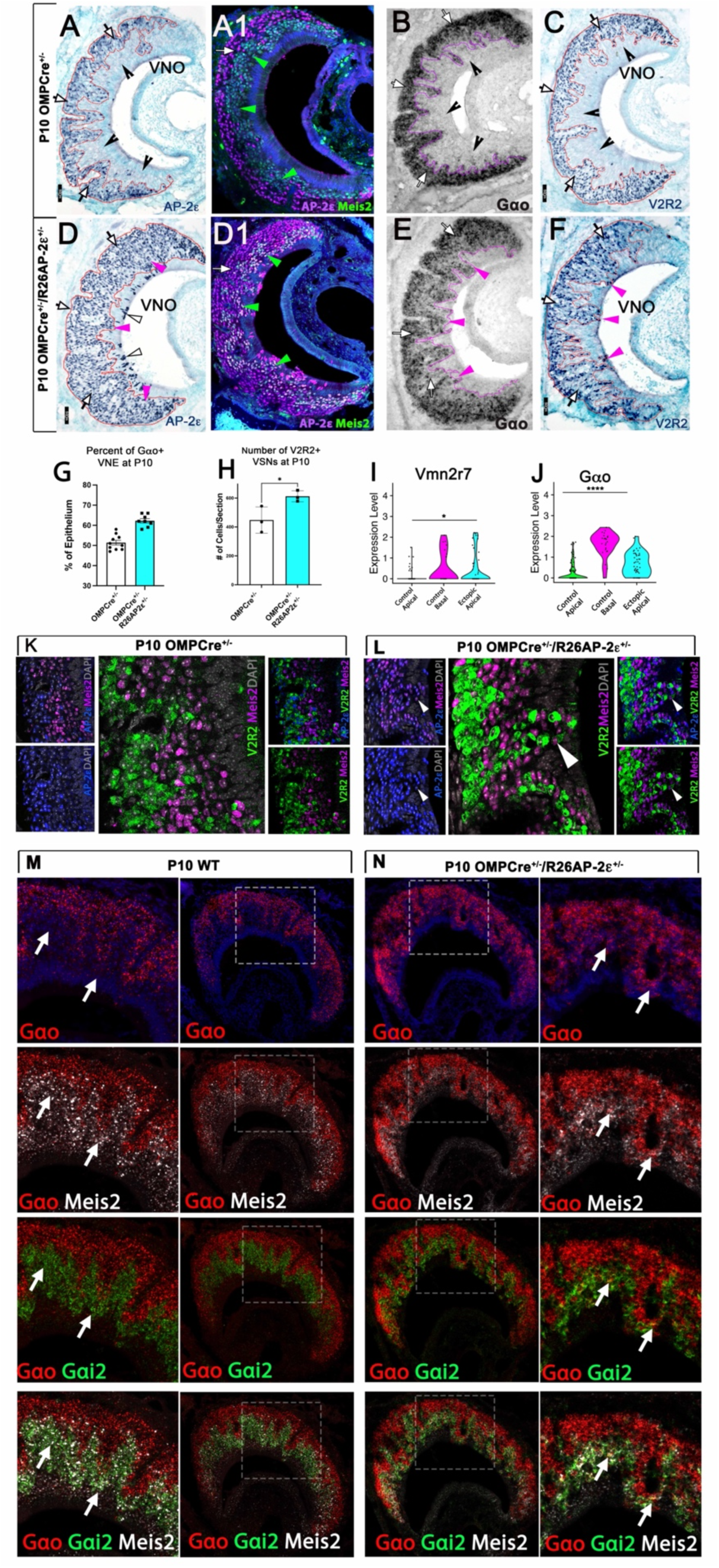
Ectopic expression of AP-2ε in the MOE and the VNE promotes expression of basal markers. Immunostainings at P10 on OMPCre^+/-^ controls (A-C) and OMPCre^+/-^/R26AP2ε^+/-^ mutants (D-F). A) IHC against AP-2ε in the VNOs of shows that AP-2ε is expressed in only the basal VSNs (arrow) in controls but have extended AP-2ε immunoreactivity into apical (magenta arrowheads) and sustentacular regions (white arrowheads) in ectopic mutants (D). A1) AP-2ε, Meis2 double immunofluorescence shows segregated AP-2ε (white arrow) and Meis2 (green arrowheads) in controls. D1) In the ectopic mutants AP-2ε expression is detected in Meis2+ cells (green arrowheads). B,E) ISH against Gαo show that in controls (B) Gαo mRNA expression is restricted to the basal regions of the VNE. E) Ectopic AP-2ε mutants show Gαo mRNA reactivity in the apical regions of the VNE (magenta arrowheads). C,F) Immunohistochemistry against V2R2 in the VNO of controls (C) shows no immunoreactivity in the apical regions of the VNE in controls (notched arrows) and is limited to the basal VSNs (white arrows). F) In mutants, more of the vomeronasal epithelium was positive for V2R2 in mutants, as expression expands into the apical regions of the epithelium (magenta arrowheads). G) Quantifications at P10 show an increase in the amount of the neuroepithelium positive for Gαo in OMPCre^+/-^/R26AP-2ε^+/-^ mutants. N=10 for OMPCre^+/-^ controls and n=8 for OMPCre^+/-^/R26AP-2ε^+/-^ mutants. H) Quantifications at P10 show a significant increase in the number of the VSNs positive for V2R2 in OMPCre^+/-^/R26AP-2ε^+/-^ mutants. N=3 for both OMPCre^+/-^ controls and OMPCre^+/-^/R26AP-2ε^+/-^ mutants (p<0.05 = *). I,J) Violin plot of Vmn2r7 (I), and Gαo (J) mRNA expression between apical and basal VSNs in OMPCre^+/-^ controls and apical VSNs of OMPCre^+/-^/R26AP2ε^+/-^ mutant mice show significant upregulation of these basal markers in mutants based on p value (p<0.05 = *, p<0.0004 = ****). K,L) K) Immunofluorescence against V2R2 (magenta) and Meis2 (green) in controls (K) and OMPCre^+/-^/R26AP2ε^+/-^ mutants (L). Arrows indicate a Meis2+ cell immunoreactive against anti V2R2 antibodies in mutants (L). M,N) Single-molecule FISH (RNAscope) against Gαo, Gαi2 and Meis2 of P10 WT and OMPCre^+/-^/R26AP2ε^+/-^ mutants. In WTs (M), a clear segregation between the Gao + (red) basal VSNs and the apical cells positive for Meis2+ (white) and Gai2+ (green) could be seen. In OMPCre^+/-^/R26AP2ε^+/-^ mutants (N) low but obvious expansion of Gao (red) expression to the apical domains of the VNO. Signal highlighting Gao expression could be found in Meis2+ (white) and Gai2+ (green) cells.

When comparing the VNO of P10 controls (either WT or OMPCre^+/-^) and OMPCre^+/-^/R26AP-2ε^+/-^ mutants, we observed that the ectopic mutants had a significantly broader Gαo mRNA expression across the VNE (Figure 4B,E). As the broader expression was hardly conducible to individual cells we performed a densitometric analysis of sections after *in-situ* hybridization (ISH). This indicated that, in the mutants, a larger percentage of the VNE was positive for Gαo expression (Figure 4G). Moreover, we found that OMPCre^+/-^/R26AP2ε^+/-^ mutants had a larger number of cells immunodetectable using the anti V2R2 antibody (Silvotti et al., 2011) (Figure 4C,F,H). These immunoreactive cells could be found spanning from basal VNO regions to the lumen.

In line with these observations scRNA-Seq data from OMPCre^+/-^ controls and OMPCre^+/-^/R26AP2ε^+/-^ indicated that apical VSNs expressing AP-2ε had variable, but significant (P<0.05) upregulation, of Gαo mRNA as well as increased expression for some C family Vmn2r, many of which can be detected using the anti V2R2 antibody (Figures 4I,J; S2E-H) (Silvotti et al., 2011). Notably our scRNA-Seq data indicated that that low/basal levels of Vmn2r7 mRNA expression can be also detected in sparse apical neurons of controls (Figures 4I; S2E-H).

Immunostaining against V2R2 and Meis2 also highlighted that, while in controls, Meis2 and V2R2 remained segregated (Figure 4K), in the ectopic mutants, V2R receptors could be immunodetected in Meis2+ apical neurons (Figure 4L). These data suggest that ectopic AP-2ε expression can induce or increase the expression of basal enriched genes.

In order to better follow the effects of ectopic AP-2ε expression on apical cells we also performed RNAscope analysis using probes against the basal VSN marker Gαo and the apical markers Gαi2 and Meis2 (Figure 4M,N).

In controls (Figure 4M), we could observe a clear segregation between the basal VSNs positive for Gαo and the apical VSNs positive for Meis2 and Gαi2. Notably Meis2 was also expressed in the sustentacular cells and in newly formed cells in the marginal zone. However, in OMPCre^+/-^/R26AP2ε^+/-^ mutants (Figure 4N), we could observe, as after regular ISH (Figure 4E) a low but obvious expansion of Gαo expression to the apical domains of the VNO. Signal highlighting Gαo expression could be found in Meis2 and Gαi2 positive cells.

### Ectopic AP-2ε expression leads to a progressive disorganization of the VNE

In ectopic AP-2ε mutants at P21, and more dramatically at 3 months of age, we noticed an increasing level of cellular disorganization of the VNE not seen in controls (Figure 5A,A’) with: 1) VSNs spanning from the basal territories to regions of the lumen devoid of Sox2+ sustentacular cells, and 2) ectopic sustentacular cells organized in spherical structures or intraepithelial cysts with a subsidiary lumen within apical and basal territories (Figure 5B-C’). Notably, the regions with ectopic sustentacular cells appeared to be mostly surrounded by apical VSNs expressing AP-2ε, Meis2, and Sox2 and were enriched in the intermediate zones of the VNE (Figure 5B-D). Interestingly, a low level of Sox2 immunoreactivity was observed in apical VSNs in both controls and OMPCre^+/-^/R26AP2ε^+/-^ mice with higher intensity in cells closer to the sustentacular cell layer (Figure S3A,B,D,E).

**Figure 5:**
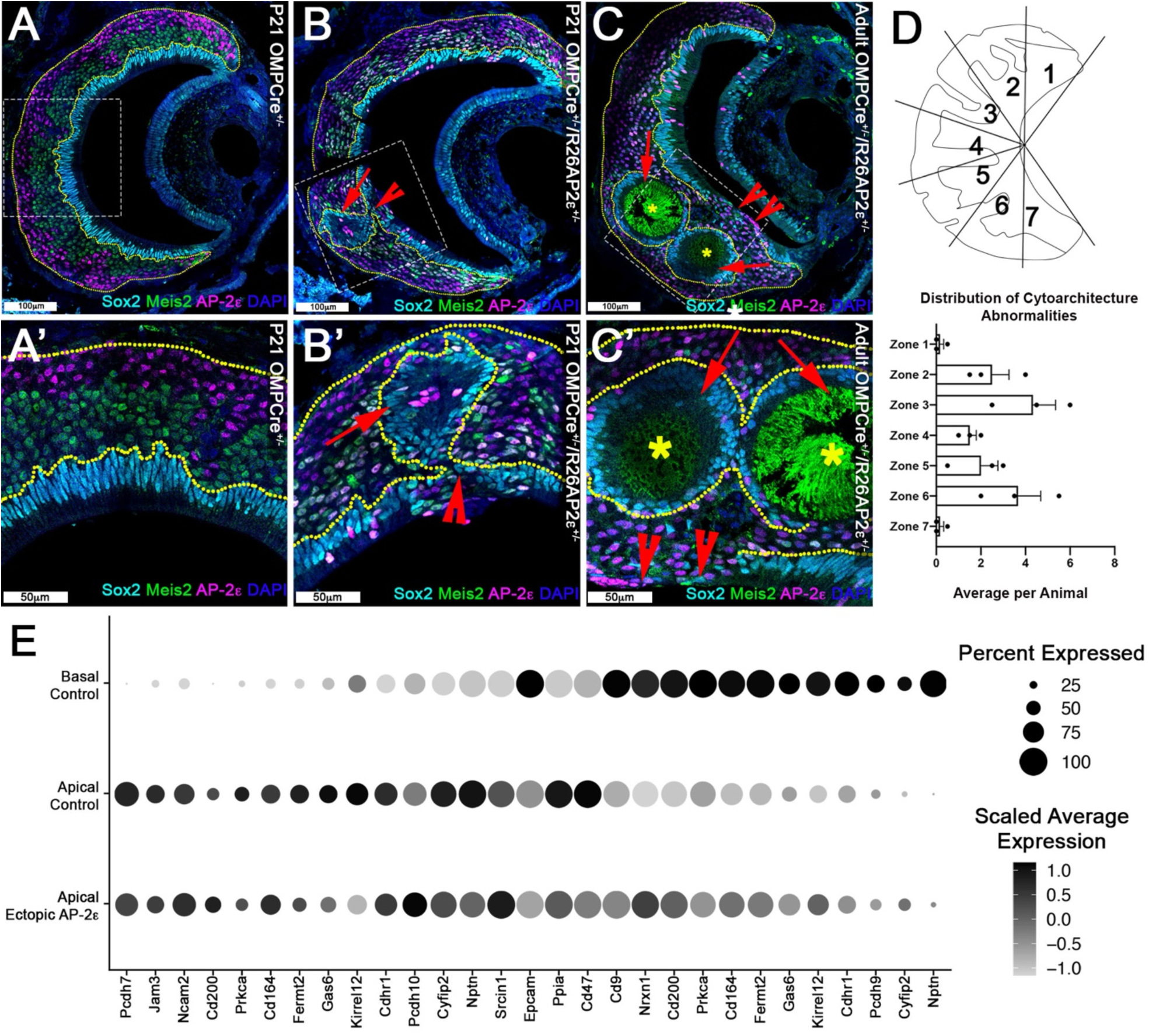
Progressive changes in VNE lamination may reflect the changing expression profiles of cell adhesion molecules in apical VSNs. Immunofluorescence against Sox2 (cyan), AP-2ε (magenta), and Meis2 (green) with DAPI (blue) counterstain. Neuroepithelium traced in yellow dotted line. A) P21 OMPCre^+/-^ controls show highly organized stratified neuroepithelium with contiguous layers of AP-2ε+/basal (magenta), Meis2+/apical (green), and Sox2+/sustentacular cell (cyan) layers. B-C’) OMPCre^+/-^/R26AP2ε^+/-^ VNO at B) P21 show that Sox2+/sustentacular cells have intraepithelial cysts with internalized subsidiary lumens (red arrows). C) Adult (3mo) OMPCre^+/-^/R26AP2ε^+/-^ mutants show an increase in the severity of intraepithelial cysts (red arrows) and breaks in the sustentacular layer and expansion of neurons to the luminal surface (red notched arrows). Unidentified matter (*) reactive to anti-mouse Abs was detected within the cysts. D) Quantifications of the zonal distribution through Zone 1 (dorsal) -> Zone 7 (ventral) of these cytoarchitecture abnormalities (which include both cell body abnormalities and dendritic disorganization, each point = 1 animal) show that these disruptions occur in the intermediate and central regions of the VNO, but not in the marginal zones (Zones 1,7). The highest rate of occurrence are in zone 3 and 6, which are intermediate regions in the VNO. N=21 E) Dot plot showing the composition and intensity of differentially expressed genes involved in cellular adhesion in the VSNs of controls and ectopic AP-2ε mutants.

The affinity and positioning of epithelial cells are largely dictated by the expression of surface adhesion molecules (Fagotto, 2014; Polanco et al., 2021). Transcriptome comparison of OMPCre^+/-^/R26AP2ε^+/-^ mutants and controls suggest that the aberrant cell positioning in the VNE of mutants can arise from broad variations in expression levels of multiple adhesion molecules throughout Meis2+ cells (Figure 5E).

Furthermore, scRNA-Seq of the adult OMPCre^+/-^/R26AP2ε^+/-^ allowed us to understand whether sustentacular cells with ectopic AP-2ε expression was contributing to the disorganization of the VNE. By performing differential gene expression analysis on the AP-2ε positive and negative sustentacular cells from the adult OMPCre^+/-^/R26AP2ε^+/-^ mice we observed significantly dysregulated genes (550 upregulated; 571 downregulated, adjusted p-value<0.05) with enrichment of genes related to tight-junctions, cell-cell adhesion, and cytoskeletal organization (Figure S3G-I), which may contribute to the disorganized neuroepithelium.

### Ectopic expression of AP-2ε alters the transcriptional profile of apical neurons

To further elucidate the gene expression changes in apical (Meis2+) VSNs after AP-2ε expression, we analyzed the UMAPs using scSeq from VSNs in controls and mutant mice. These revealed similar clustering at the stages of neurogenesis and differentiation across genotypes (Figure 7A-B). However, control animals showed AP-2ε expression was limited to immature-mature basal VSNs (Figures 7A’-A’’’; 4A1). In OMPCre^+/-^/R26AP-2ε^+/-^ mutants, however, AP-2ε mRNA was expressed in maturing basal VSNs as well as in maturing and mature apical VSNs (Figures 7B’, B’’’; 4D1). When analyzing the UMAPs, we noticed that in the OMPCre^+/-^/R26AP-2ε^+/-^ mutants, cells along the apical and basal developmental trajectories overlapped to those of controls. However, the maturing and mature apical VSNs of controls and mutants formed non overlapping clusters (Figure 7A’’’-(A+B)’’’’). In fact, the apical VSNs of the ectopic mutants formed a cluster more proximal to the basal VSNs.

**Figure 7:**
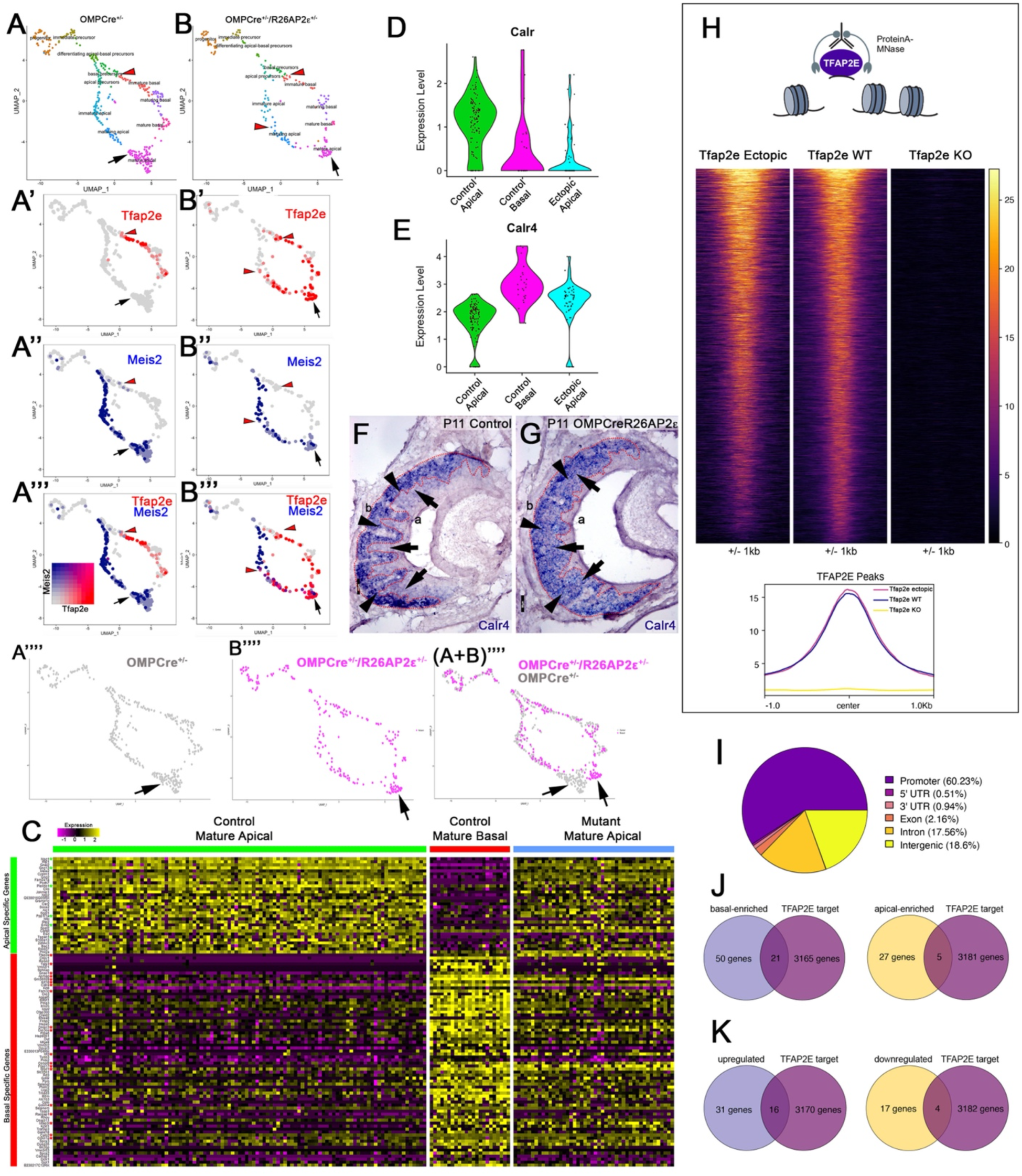
Single-cell sequencing of P10 OMPCre^+/-^ control and OMPCre^+/-^R26AP2ε^+/-^ mutant VSNs indicate a shift in apical cells towards basal cells in the mutant. A-A’’’) UMAP clustering of VSNs from progenitor cells to differentiated mature apical and basal cells of Control B-B’’’) Mutant mice split by genotype. A’-A’’’, B’-B’’’) Blended feature plots of AP-2ε expression (red) and Meis2 (blue). Red arrowheads indicate onset of AP-2ε expression. Black arrow indicates mature apical VSNs. OMPCre^+/-^ Controls (A’-A’’’) show a divergent pattern of expression where the onset of AP-2ε (red, red arrowhead) is only on the basal branch. Meis2 expression (blue) occurs only on the apical branch where the cells lack AP-2ε expression. OMPCre^+/-^R26AP2e^+/-^ mutants (B’-B’’’) start to express AP-2ε on the basal branch in immature basal VSNs (red) however, onset of AP-2ε mRNA expression also occurs on the apical branch (blue). B’’’) In ectopic mutants, AP-2ε mRNA is co-expressed with Meis2 in apical VSNs (purple cells). A’’’’) Feature plot of apical cells in OMPCre^+/-^ controls (black arrow). B’’’’) Feature plot of apical cells in OMPCre^+/-^R26AP2e^+/-^ mutants (black arrow). (A+B)’’’’) Overlay shows that apical cells with ectopic AP-2ε expression (magenta, black arrow) clustered separately from mature apical cells of OMPCre^+/-^ controls (gray, black arrow). D) Violin plot shows Calreticulin (Calr) mRNA expression levels in apical VSNs from the OMPCre^+/-^R26AP2ε^+/-^ mutants are reduced to levels similar to basal VSNs from OMPCre^+/-^ controls. E) Violin plot shows Calreticulin-4 (Calr4) mRNA expression levels. E) In control cells, AP-2ε mRNA expression (red arrowheads) and Meis2 mRNA expression (blue) are not co-expressed in the same cells. AP-2ε expression is upregulated in immature basal VSNs and not apical VSNs. F,G) ISH against Calr4 against P11 OMPCre^+/-^ controls (F) and OMPCre^+/-^R26AP2ε^+/-^ mutants (G) show that while Calr4 mRNA is normally enriched in basal VSNs (arrowheads), ectopic AP-2ε mutants show expansion of Calr4 positivity in the apical regions when compared to controls (arrows). H-K) Analysis of CUT&RUN against Tfap2e/AP-2ε. H) Tornado plot of AP-2ε occupancy in Tfap2e ectopic, WT and KOs in the dissociated tissue of the VNO. Tfap2e signal in ectopic mutants and WTs is similar across all Tfap2e peaks, while Tfap2e KOs show no signal. The genomic regions are defined as the summit +/-1kb. I) Pie chart depicting the genomic distribution of putative AP-2ε binding sites show that most of AP-2ε peaks are found in promoter regions of putative target genes and to a lesser extent in intergenic and intronic regions of the genome. J) Venn diagram of the determined AP-2ε targets and the genes enriched in the basal and apical VSNs. K) Venn diagram of the determined AP-2ε targets and all the upregulated and downregulated genes in the ectopic AP-2ε mutant mouse. Significance defined as Adjusted p-value ≤ 0.05.

To understand the extent to which AP-2ε can reprogram apical VSNs, we further compared the expression of the most enriched genes in apical and basal VSNs of OMPCre^+/-^ controls to the apical VSNs of OMPCre^+/-^/R26AP-2ε^+/-^. Interestingly, this analysis revealed that apical VSNs of OMPCre^+/-^/R26AP-2ε^+/-^ mice had a mixed apical-basal RNA expression profile with a significant down-regulation of ∼22% of the apical-enriched genes (7/32), and a significant upregulation of ∼28% of the basal-enriched genes (20/71) (Figure 7C; Supplementary Table S2). Performing a correlation analysis, we observed that while in controls, sets of either apical or basal enriched genes had high correlation, this was no longer true for the OMPCre^+/-^/R26AP-2ε^+/-^ mutants (Figure S4).

Of the aberrantly expressed genes in the apical VSNs of OMPCre^+/-^/R26AP-2ε^+/-^ mice, we identified a reduction in Calreticulin (Calr) mRNA levels together with a strong upregulation of Calreticulin4 (Calr4), which persists in adulthood (Figure 7D-G, Supplementary Table S2, Figure S8A,C). Calr is a negative regulator of transport of V2R receptors to the cell membrane (Dey and Matsunami, 2011). ISH at P11 confirmed that Calr4 is normally expressed by basal VSNs in controls. However, in OMPCre^+/-^/R26AP2ε^+/-^ mutants, Calr4 mRNA was found in both apical and basal VSNs (Figure 7F,G). In line with previous studies (Dey and Matsunami, 2011), we found that Calr was expressed below ISH detectability. Feature maps also pointed to the ectopic expression in apical VSNs of other basal enriched genes such as Gαo (Figures 4; S5A), Mt3, and Fbxo17 (Figure S5B,D). In apical cells these genes are normally either silenced in mature VSNs or absent from the beginning of the differentiation. RNAscope analysis for Mt3 confirmed ectopic expression in apical cells (Figure S5C).

### OMPCre^+/-^/R26AP-2ε^+/-^ ectopic mutants have normal axonal projections to the AOB

Axonal projection along the anterior posterior axis of the AOB is largely determined by axon guidance molecules such as Nrp2, Robo2 while the the coalescence of vomeronasal sensory neuron axons into glomeruli is largely dictated by Kirrel adhesion molecules (Cloutier et al., 2002; Prince et al., 2013; Prince et al., 2009; Vaddadi et al., 2019). The mRNA expression levels for the guidance receptors, Robo2 and Nrp2, and the adhesion molecules, Kirrel2 and Kirrel3, did not significantly change after ectopic AP-2ε expression. In fact, by immunostaining against Robo2 and Nrp2 we confirmed immunoreactivity of Nrp2 in the anterior and Robo2 in the pAOB similar to controls and observed no significant differences in the average size of anterior or posterior AOB between genotypes (Figure S6 A,B,E). Moreover, quantifications based on Kirrel2 and Kirrel3 immunostaining did not reveal major changes in glomerular size or number in the AOB (Figure S6C,D,F,G) (Bahreini Jangjoo et al., 2021).

### Ps6 immunostaining reveals that OMPCre^+/-^/R26AP-2ε^+/-^ mutants have defective response to female urines

Whole male mouse urine activates both V1Rs and V2Rs (Krieger et al., 1999), while female odorants mostly activate apical VSNs (Dudley and Moss, 1999; Kimoto et al., 2005; Norlin et al., 2001; Silvotti et al., 2018). To determine if OMPCre^+/-/^R26AP-2ε^+/-^ mice had altered chemodetection, we quantified VSNs’ activation after exposure of control and mutant mice to either male or female soiled bedding. Brains were collected after 90 minutes of exposure to the soiled bedding, to allow adequate time for the phosphorylation of the ribosomal protein S6 (pS6) in the VSNs’ cell bodies (Silvotti et al., 2018). VSNs activation was quantified after immunostaining against pS6 (Ser 240/244) on coronal sections of the VNO (Figure 6A-D). Apical and basal VSNs were identified with immunostaining against Meis2 and categorized as either pS6+/Meis2+ apical or pS6+/Meis2-basal VSNs (Figure 6A-D). When exposed to male bedding we observed that OMPCre^+/-^/R26AP-2ε^+/-^ mice had a lower average number of activated apical VSNs compared to controls, however this difference was non statistically significant (Figure 6A,B,E). Although this difference was non statistically significant, we did find a significant reduction in the total activation of VSNs in female mutants.

**Figure 6:**
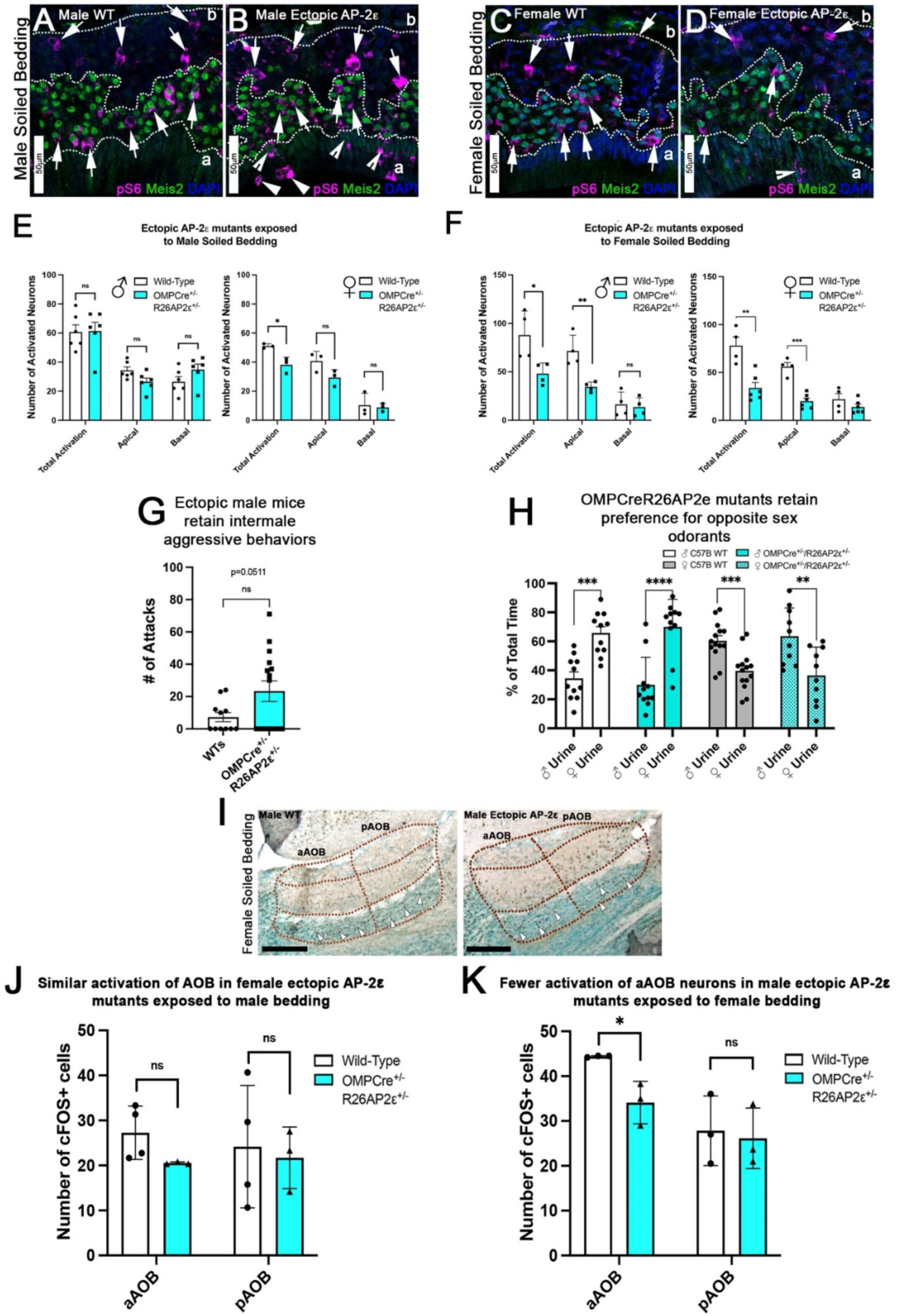
Ectopic AP-2ε expression alters the detection of sex-specific odorants. A-D) Immunofluorescence against pS6 (magenta, arrows) and Meis2 (green) with DAPI counterstain (blue) in the VNE of Controls (A,C) and OMPCreR26AP-2ε mutants (B,D). A,B) VNE of adult male wildtype and ectopic mutants when exposed to male soiled bedding show similar activation or pS6 immunoreactivity (arrows) in Meis2+/apical and Meis2-/basal VSNs. C,D) VNE of adult female wildtype and ectopic mutants when exposed to female soiled bedding show that while WT females displayed a higher proportion of pS6+ apical VSNs, OMPCre^+/-^/R26AP-2ε^+/-^ mutants showed a decreased number of activated Meis2+/apical VSNs. In both exposure conditions ectopic activation of sustentacular cells (notched arrows) and VSNs near the lumen (arrowheads). E) Quantifications of activated VSNs in male and female mice after exposure to male-soiled bedding show a non-significant decrease in activated apical VSNs in mutants of both sexes. Mutant females show a significant decrease in total activation of VSNs (p<0.05) while mutant males show a small but non-significant increase in the total number of activated basal VSNs. N=6 for males and n=3 females for both genotypes F) Quantifications of activated VSNs after exposure to female-soiled bedding in male and female mice show a significant decrease in the number of activated apical VSNs as well as total activation of VSNs between mutants and controls of both sexes (p<0.05=*, p<0.01=**, p<0.001=***). N=4 for males and n=6 for females for both genotypes. G) Quantifications of the number of attacks for all WT and OMPCre^+/-^/R26AP-2ε^+/-^ mutants in a resident intruder test show male ectopic AP-2ε mice display higher levels of intermale aggression but are not significantly different than controls. N=11 for WTs and n=17 for OMPCre^+/-^/R26AP-2ε^+/-^ mutants. H) Quantifications of odorant preference tests in male and female mice show that male and female OMPCre^+/-^/R26AP-2ε^+/-^ mutants retain the preference for opposite sex odorants. (p<0.05=*, p<0.01=**, p<0.001=***, p<0.0001=****). N=11 for WT males, n=14 for WT females, n=11 for OMPCre^+/-^/R26AP-2ε^+/-^ mutant males and n=10 for OMPCre^+/-^/R26AP-2ε^+/-^ mutant females. I) AOB of adult male wildtype and ectopic mutants when exposed to female soiled bedding show that ectopic mutants have lower levels of cFOS activated neurons in the anterior AOB (aAOB) but similar levels of activated neurons in the posterior AOB (pAOB). Scale bar=250um. J) Quantification of cFOS activated neurons in the AOB of female mice exposed to male-bedding show no significant differences between anterior and posterior AOB activation in mutants and WT. N=4 for WT and n=3 for OMPCre^+/-^/R26AP-2ε^+/-^ mutants. K) Quantification of cFOS activated neurons in the AOB of male mice exposed to female-soiled bedding show a significant reduction in anterior AOB activation (p<0.05) in mutants compared to WT. N=3 for both WT and OMPCre^+/-^/R26AP-2ε^+/-^ mutants.

In order to further analyze the activity of apical VSNs we exposed control and OMPCre^+/-^/R26AP-2ε^+/-^ males and females to female soiled bedding which mostly activates apical VSNs. This experiment highlighted a dramatic reduction in apical VSNs’ activation, as well as total activation of VSNs (pS6+/Meis2+) of mutant mice for both sexes (Figure 6C,D,F).

### Ectopic AP-2ε enhances intermale aggressive behavior but not preference for opposite sex odorants

To determine whether the aberrant gene expression in apical VSNs could alter VSNs’ functions and related social behaviors, we evaluated intermale aggression and odorant preference (Koolhaas et al., 2013). By performing resident intruder tests, we showed that the level of intermale aggression of OMPCre^+/-^/R26AP-2ε^+/-^ male mice was higher but not significantly different than controls (P=0.051) (Figure 6G). However, among the animals that displayed aggressive behavior we observed significant increase in the number of attacks from WT to OMPCre^+/-^/R26AP-2ε^+/-^ mutants (p=0.0015; WT=16.00 SE+/- 3.1; Ectopic=43.75 SE+/- 4.8).

The sex urines preference test revealed that both male and female OMPCre^+/-^/R26AP-2ε^+/-^ mutants exhibited preferential interest in opposite sex urines similar to controls (Figure 6H). Interestingly, OMPCre^+/-^/R26AP-2ε^+/-^ females, displayed much more variability in their individual odor preferences compared to controls, nonetheless the ectopic mutants still retained their preference for opposite sex odorants.

All together these data suggest that ectopic AP-2ε expression decreases apical VSNs’ functionality (Figure 6A-F) likely leading to an increase in aggression behavior but not compromising opposite sex odorants preferences (Figure 6G,H). These data are in line with previous findings indicating that loss of apical VSN signal transduction enhances territorial aggression in males without substantial changes in sex odor preferences (Trouillet et al., 2019).

### The negative effects of ectopic AP-2ε expression in apical neurons functionality is reflected by reduced c-Fos activation in the anterior AOB

To further investigate if AP-2ε ectopic expression in apical neurons alters the vomeronasal signal transduction, we analyzed c-Fos activation in the AOB. To do this we analyzed the AOBs of control and OMPCre^+/-^/R26AP-2ε^+/-^ animals exposed to opposite sex soiled bedding. This analysis revealed that c-Fos activation in the aAOB was statistically different only after exposure to female soiled bedding (Figure 6I,K), while there was a non-significant reduction in c-Fos activation in the aAOB of female OMPCre^+/-^/R26AP-2ε^+/-^ mice exposed to male bedding when compared to wildtypes (Figure 6J). These data suggest that ectopic AP-2ε expression reduces the functionality of the V1R VSNs projecting to the aAOB but does not alter the functionality of basal VSNs.

### Identification of direct AP-2ε targets via CUT&RUN

Our findings suggest a key role for AP-2ε in controlling the expression of specific basal specific/enriched genes. Transcriptomic studies in AP-2ε^Null^ mice showed loss of expression of basal VSN specific genes suggesting that AP-2ε controls parts of the basal and apical VSN genetic programs (Lin et al., 2018). However, it remains unknown whether AP-2ε regulates VSN genetic programs directly or indirectly. So, we performed genome-wide mapping of transcription factor occupancy with cleavage under targets and release using nuclease (CUT&RUN) to determine the direct genetic targets of AP-2ε in the VNO to pair with our scRNA-Seq (Figure 7H-K). Our analyses identified over 5025 replicable peaks in VNO tissue indicating AP-2ε binding sites. Notably, performing CUT&RUN and sequencing from AP-2ε KOs revealed 203 peaks, of which 154 overlapped with called WT peaks. After subtracting out peaks called in the knockout, we were left with 4871 peaks (Figure 7H). AP-2e peaks of the WT were assigned to 3186 genes. None of the peaks of the AP-2ε KO that were subtracted out were associated with apically or basally enriched genes. CUT&RUN peaks of ectopic mutants were largely similar to that of WTs (Figure 7H).

Of these putative binding sites, we found that 60.23% of the peaks occurred in promoter regions (defined as any region 1000bp upstream or 200bp downstream a transcription start site), 18.6% in distal intergenic regions, and ∼17.56% in intronic regions (Figure 7I). These results suggest that most AP-2ε activity directly regulates transcription, with perhaps a secondary role in enhancer regions.

Gene Ontology analysis of genes associated with AP-2ε peaks showed an enrichment of factors involved in protein degradation, transcription coregulator activity, and histone and chromatin modification (Figure S7A). Motif enrichment analysis of AP-2ε peaks revealed Tfap2 as the top enriched motif (p=1e-180), as expected. Other transcription factor motifs enriched in the same regions as AP-2ε peaks include SP, KLF, EBF, RFX, NRF, DLX, and LHX transcription factor families (Figure S7B). As many transcription factors work with other cofactors to regulate gene expression (Huang et al., 2015; Monahan et al., 2017), these motifs may represent potential cofactors that work in concert with AP-2ε to mediate either an activating or repressive role in the VNO.

When we compared AP-2ε direct targets with our identified apical and basal enriched genes from mature VSN populations, we discovered that 18% of our identified apical-enriched genes (5/27 genes) (Supplementary Table 1) were AP-2ε direct targets and approximately 47% of our identified basal-enriched genes (21/50 genes) are AP-2ε direct targets (Figure 7J). Of the most canonical apical and basal markers and signal transduction machinery only Gαi2/Gnai2 had putative direct AP-2ε occupancy and assumed regulation of transcription (Figure S7C). Out of our newly discovered list of basal enriched genes (Supplementary Table 1) we identified that Krt18 has a putative AP-2ε binding site within its promoter region. As expected from a terminal selector gene, these data suggest that AP-2ε directly binds and regulates batteries of apical and basal enriched genes (Figure S7C).

These data suggest that AP-2ε plays a dual role in maintaining the basal VSNs’ genetic program while restricting the expression of genes normally enriched in apical VSNs (Figures 7J,K; S7). In line with this, CUT&RUN from ectopic expressors gave tracks that largely overlapped with those of the WT controls (Figure S7) (Figure 7H). However, when plotting the signals at the promoter of apical and basal enriched genes for WT, AP-2ε KO and AP-2ε ectopic expressors we observed more signal at the apical-enriched promoters in the ectopic mice dataset (Figure S7D). This data suggests that ectopic expression of AP-2ε in apical neurons facilitates its access to the promoter of apical enriched genes as these are normally active in the apical neurons.

## DISCUSSION

Understanding how differentiated neurons retain cellular plasticity remains critical to identify how genetic insults can compromise neuronal identity, circuit assembly and function (Hobert and Kratsios, 2019; Molyneaux et al., 2007; Patel and Hobert, 2017; Pereira et al., 2019; Rahe and Hobert, 2019). Spatial and temporal expression of terminal selector genes regulates the establishment and maintenance of neuronal identity remains foundational to elucidate the assembly of functional neuronal circuits (Arlotta et al., 2005; Cau et al., 2002; Cau et al., 1997; Molyneaux et al., 2007). In fact, loss of terminal selector genes can lead to loss of neuronal identity and increase cellular/phenotypic plasticity, while expression of specific TFs can induce specific cellular features only at particular developmental windows (Hobert, 2008; Rahe and Hobert, 2019).

Rodents and some marsupials have a binary vomeronasal epithelium where the two main types of VSNs, apical and basal VSNs, are generated throughout life from a common pool of Ascl1 progenitors (Berghard and Buck, 1996; Jia and Halpern, 1996; Katreddi and Forni, 2021; Mohrhardt et al., 2018; Silva and Antunes, 2017; Taroc et al., 2020; Weiler et al., 1999). The generation of these two distinct populations is central for critical socio-sexual behavior in rodents (Oboti et al., 2014; Perez-Gomez et al., 2014). In a recent study we have shown that the apical-basal differentiation dichotomy of VSNs is dictated by Notch signaling. The transcription factor TF AP-2ε is expressed in maturing cells fated to become V2R neurons. In this study, we combined scRNA-Seq, histology, behavior, and CUT&RUN methodologies to test if TF AP-2ε is a basal VSN specific terminal selector gene capable of partially reprogramming the apical VSN identity.

Using scRNA-Seq, we discovered key transcriptomic differences between mature basal and apical VSNs that were previously unreported (Figure 1Q, Supplementary Table S1). We also confirmed that AP-2ε mRNA is restricted to maturing Gαo/basal VSNs (Figure 1K) and that AP-2ε itself does not initiate the basal VSN differentiation program, but rather maintains the integrity of the basal neuronal identity. In fact by re-expressing AP-2ε in AP-2ε KOs we demonstrated that AP-2ε is indispensable for basal cellular homeostasis (Figure 2) and therefore for the establishment of normal territorial and sex-preference behaviors of rodents (Figure 3). We also elucidated that AP-2ε acts in controlling VSN gene expression through activating and repressive activity when analyzing mature/maturing Meis2+ apical VSNs in OMPCre^+/-^/R26AP-2ε^+/-^ mutant mice (Figure 7).

During differentiation chromatin barriers dynamically restrict the cellular plasticity, preventing ectopic terminal selector genes from genetically reassigning neurons (Rahe and Hobert, 2019). We have previously shown that postmitotic VSNs of AP-2ε KO mice can partially deviate from the basal differentiation program and turn on sets of apical specific genes. However, AP-2ε LOF did not prevent basal neurons from acquiring basal features, such as Gαo or V2Rs ((Lin et al., 2018), suggesting that AP-2ε activity is crucial to restrict basal VSN phenotypic plasticity rather than establishing the basal cell fate (Lin et al., 2018). Here we showed that AP-2ε null mice have reduced odorant sex preference and intermale aggressive behavior, which are classic phenotypes related to basal VSN LOF (Stowers et al., 2002). However, we found that reintroducing AP-2ε in maturing basal AP-2ε KO neurons was sufficient to rescue cellular homeostasis (Figure 2), physiological functions, and related behavior (Figure 3).

Terminal selectors define neuronal identity by suppressing alternative programs and can also act as pioneer factors (Lupien et al., 2008; Magnani et al., 2011; Mangale et al., 2008). Based on our rescue data, we propose that AP-2ε can partially reprogram/alter the transcriptome of differentiated cells, as expected from a pioneer factor and other members of the Tfap2 family (Rothstein and Simoes-Costa, 2020).

When we used OMPCre drivers to induce ectopic AP-2ε in differentiated olfactory and both apical and basal vomeronasal neurons, we observed progressive gene expression, and morphological changes in the VNO, but no gross morphological changes in the main olfactory epithelium (Figures 4; 5; S2). We suspect the lack of phenotype in the OE may arise from the absence of necessary Tfap2 cofactors, that are expressed in the VNO (Figure S1) (Eckert et al., 2005).

The co-expression of AP-2ε in Meis2+ apical VSNs in OMPCre^+/-/^R26AP-2ε^+/-^ mice revealed that Meis2, and most apical specific genes, were expressed at P10 (Figure 7C). However, single-cell transcriptome analyses in adult AP-2ε ectopic mice indicated that several apical genes, including Meis2, were expressed at significantly lower levels than controls (Figure S8A,C). These data suggest that AP-2ε can negatively modulate genes enriched in the apical program.

Ectopic expression of individual terminal selector genes can selectively control specific molecular features linked to neuronal function and identity, but not pan-neuronal features like guidance cue receptors (Patel and Hobert, 2017; Stefanakis et al., 2015). Our ScRNA-Seq revealed that AP-2ε ectopic expression, does not alter the expression of VSN-specific guidance cue receptors Nrp2 and Robo2 (Cho et al., 2011; Cloutier et al., 2002; Prince et al., 2009; Walz et al., 2002). Notably Nrp2 and Robo2 start to be expressed soon after the apical/basal VSNs’ developmental trajectories are established, suggesting that these genes are expressed before and independently of tfap2e expression (Figures 1; 7; S6). In line with this, we observed that AP-2ε ectopic expression did not significantly change Kirrel2 and Kirrel3 expression patterns (S6). As a result we found not significant changes in glomeruli size or number in the AOB (S6C,D,F,G).

In OMPCre^+/-^/R26AP-2ε^+/-^ mutants, we observed that the cellular organization and lamination of the vomeronasal epithelium became severely disrupted between P10 and adult ages (Figure 5). In fact, in the mutants, we found basal neurons located at the level of the VNE lumen and ectopic sustentacular cells forming intraepithelial cyst-like structures in both apical and basal territories (Figure 5B,C). Notably, the disorganization of the VNE that resembles intraepithelial cysts as previously described in aging mice (Wilson and Raisman, 1980), appeared to be more pronounced/frequent in regions proximal to the neurogenic marginal zones (Figure 5D,E), where OMP mRNA is expressed following Gap43 expression. Therefore, we posit that the regionalization of the VNE phenotypes might represent cells that underwent AP-2ε ectopic expression at early maturation stages. When we compared mRNA of control and ectopic AP-2ε mutants, which have disorganized VNE, we observed changes in expression levels of many surface and cell adhesion related molecules as well as upregulation of stress related genes (Figures 5E; S3E). In addition to this, scRNA-seq analysis of AP-2ε positive and negative sustentacular cells in adult OMPCre^+/-^/R26AP-2ε^+/-^ mutants revealed massive changes in gene expression in these support cells. Future studies should focus on understanding which of the dysregulated genes in VSNs and sustentacular cells contribute to the cytoarchitectural organization of VSNs and sustentacular cells.

Sex odorants activate different sets of vomeronasal receptors and therefore different populations of VSNs (Dudley and Moss, 1999; Keller et al., 2006; Silvotti et al., 2018). Interestingly, scRNA-seq analysis and validation via RNA scope and ISH revealed that ectopic AP-2ε in mature apical VSNs leads to the upregulation of basal enriched genes such as Gao (Figure 4) and the ER chaperone protein Calreticulin-4 (Calr4) (Figure 7). In the ectopic AP-2ε mutants we observed a reduction in mRNA of Calr (Figure 7D). Loss of Calr expression has been previously shown to increase V2R cell surface expression (Dey and Matsunami, 2011). Changes in Calr4 and Calr expression levels could be partially responsible for the overall increase in family-C V2Rs cell surface expression/immunoreactivity (Dey and Matsunami, 2011) observed in ectopic AP-2ε mutants (Figure 4C,F,H).

After OMPCre driven ectopic AP-2ε expression we observed a significant upregulation of mRNAs of V2r receptors coded by genes belonging to the C-family (Silvotti et al., 2011) (Figure S2). Notably, our data suggest that some apical neurons of controls can express low mRNA levels of family-C Vnm2r mRNAs (Figures 4; S2E-H). These data suggest that some family-C genes (e.g. Vmn2r7), which are known to not follow rigorous mechanisms of monogenic expression, might also have a much looser cell type specific expression control that previously postulated. However, using the anti V2R2 antibody, which recognized a large spectrum of Vmn2r of the family-C, including Vmn2r7 (Silvotti et al., 2011), we could detect cells immunoreactive for both V2R2 and Meis2 in AP-2ε ectopic mutants (Figure 4L). This suggests that in control animals apical VSNs either express V2R genes below immune-detectability (Fig 4K) or that post transcriptional mechanisms may play roles in silencing translation.

Using pS6 immunoreactivity to detect activated VSNs, and cFos to detect signal transduction in the AOB revealed that ectopic AP-2ε mutants have an overall decreased apical cells’ ability to detect and transduce signal to the aAOB (Figure 6). Notably these defects resulted more obvious, in both males and females, after female bedding exposure (Figure 6). Conditional ablation of normal apical VSN signal transduction does not undermine intermale aggression, rather it enhances territorial aggression in mutant males (Trouillet et al., 2019). In line with this, we found that OMPCre^+/-/^R26AP-2ε^+/-^ mutants that display aggressive behavior had higher levels of aggression towards male intruders when compared to WT controls (Figure 6G). These data indicate that ectopic AP-2ε expression in V1R neurons is sufficient to partially subvert their function and therefore alter intrinsic social behaviors.

In rodents, olfactory sex discrimination persists after VNO excision; however, preference for opposite sex odorants is mediated by the accessory olfactory system (Keller et al., 2006; Pankevich et al., 2004). Despite the partial desensitization of the apical VSNs in OMPCre^+/-/^R26AP-2ε^+/-^ mutants, transcriptome, and morphological changes in the VNO did not compromise normal sex odorant preference in male or female mutants. In fact, conditional ablation of Gαi2, which is required for normal signaling of apical VSNs, did not alter normal male sexual behavior, including male preference for estrous female urine (Trouillet et al., 2019). Our data support a dispensable role for apical VSNs neurons in the tested sexual behavior behaviors (Figure 6C,D,H).

To elucidate the mechanism of action of AP-2ε, we performed CUT&RUN (Skene et al., 2018; Skene and Henikoff, 2017). This analysis (Figure 7) identified putative direct targets of AP-2ε and showed 3000+ putative binding sites in the vomeronasal tissue. The specificity of binding was confirmed by CUT&RUN experiments on AP-2ε Kos (Figure 7H). In the identified AP-2ε target genes we found that most of binding sites were primarily in promoter regions, not in intergenic and intronic regions. These data suggest that AP-2ε’s main mechanism of action directly regulates gene transcription with perhaps a secondary role at enhancer regions. We found AP-2ε bound to the up and down-regulated genes in both apical VSNs and sustentacular cells in the OMPCre^+/-^/R26AP-2ε^+/-^ ectopic mouse line, which suggests that AP-2ε can act as both a transcriptional activator and repressor. Motif analysis of these up and downregulated regions indicates that AP-2ε may function in concert with specific transcriptional cofactors to fulfill a dual role in maintaining the basal VSN genetic program and restricting cellular plasticity (Figure S7). Notably our CUT&RUN data showed an enrichment of factors involved in transcription coregulator activity, histone modification, and chromatin modification (Figure S7A). These data indicate that AP-2ε may play a role in modifying the chromatin landscape indirectly or in tandem with these transcriptional cofactors to regulate the basal genetic program. Even though transcriptome and histological analysis of the VNE showed significant changes in canonical apical and basal specific genes, we only Gαi2 and Krt18 had AP-2ε peak assignments, suggesting that AP-2ε acts indirectly to regulate these genes. However, since peaks were assigned to the nearest gene, we cannot exclude long-distance gene regulation through enhancer regions as a contributor. CUT&RUN peaks of ectopic mutants were largely similar to that of WTs, while only background peaks were found in AP-2ε KOs. Notably, our data suggests that ectopic expression of AP-2ε in apical neurons allows for access of Tfap2e to the promoter of apical enriched genes as they are in the euchromatic state (S7).

In conclusion, the results of our study indicate that AP-2ε has some features of a terminal selector gene. In fact, AP-2ε plays roles in controlling the expression or expression levels of several basal enriched VSN genes and repressing apical ones, which is necessary for normal basal VSN functions that mediate territorial and sex preference behaviors in mice. We recently found that the establishment of the apical/basal identity is a slow and multistep process, and that the apical/basal identity is primarily established by Notch signaling as soon as the cells become postmitotic (Katreddi et al., 2022). In line with this, we observed that after ectopic AP-2ε expression in maturing apical neurons the core of their default apical identity is maintained, however some of the genes normally expressed in alternative basal identity are turned on in apical neurons. Our data suggest that after AP-2ε expression the apical VSNs acquire an ambiguous transcriptome identity as ectopic AP-2ε expression is sufficient to bypass some layers of cellular plasticity restrictions over time. These changes translate into a reduced functionality of the apical neurons, rather than into a transdifferentiation to basal VSNs. The genetic changes induced by ectopic AP-2ε further manifest in a progressive disorganization of the vomeronasal neuroepithelium like that reported in aging animals (Wilson and Raisman, 1980).

Our study suggests that as previously hypothesized by others (Hobert and Kratsios, 2019; Rahe and Hobert, 2019) aberrant expression of terminal selector genes in postnatal neurons can alter the transcriptomic identity of neurons, organization neuroepithelia, and potentially lead to neuropathologies.

## STAR**★**METHODS

### Key Resources Table

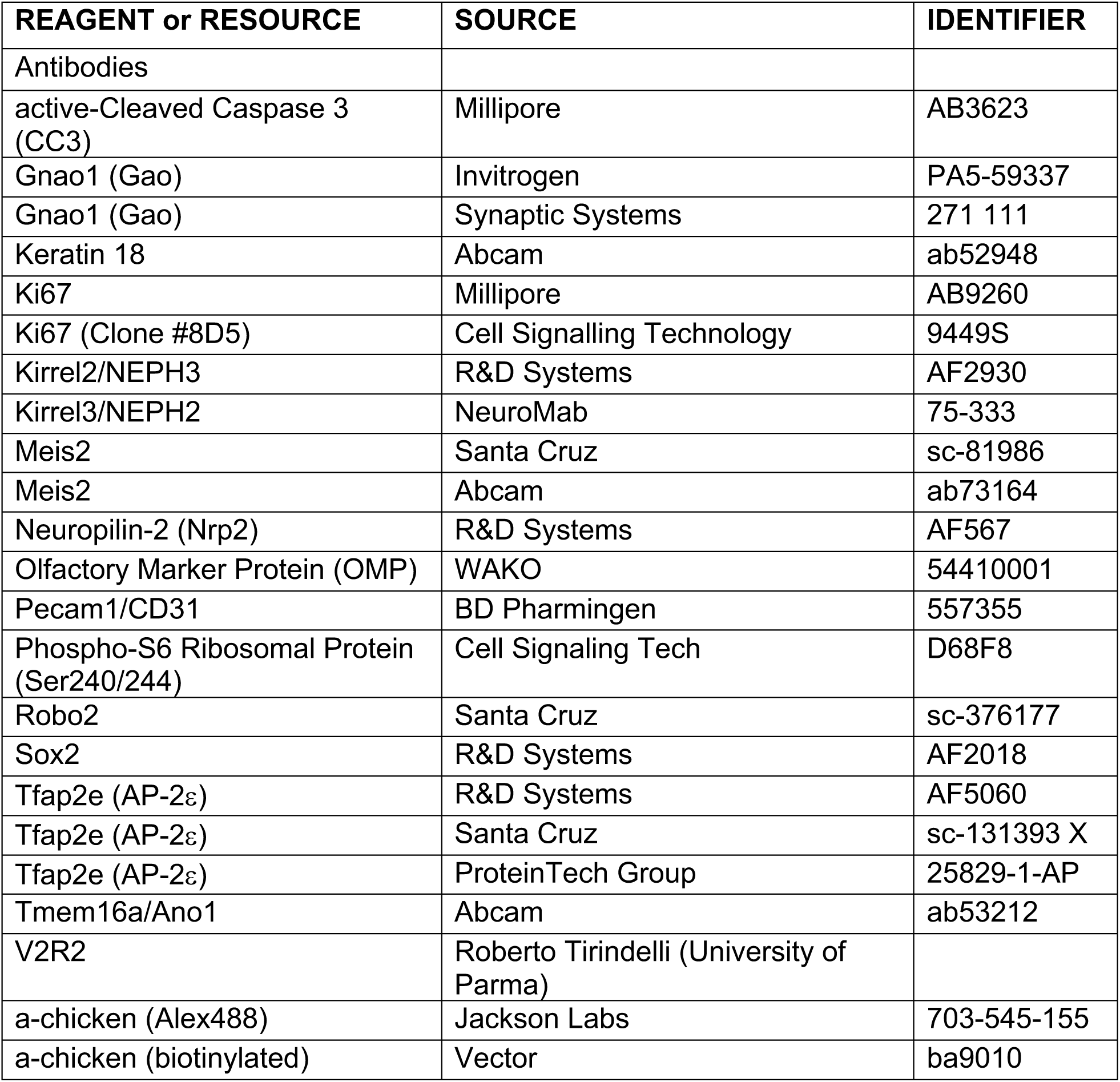
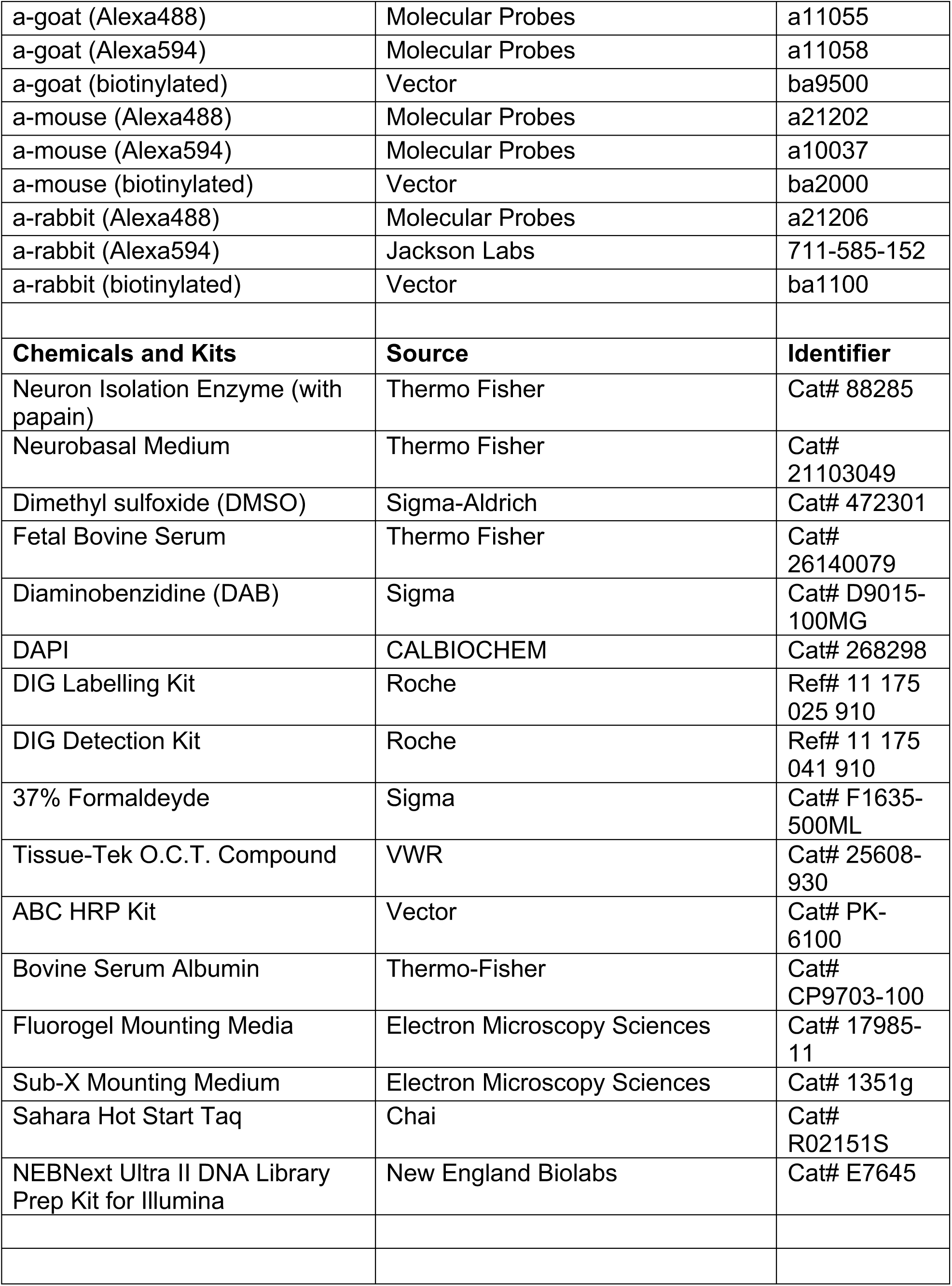
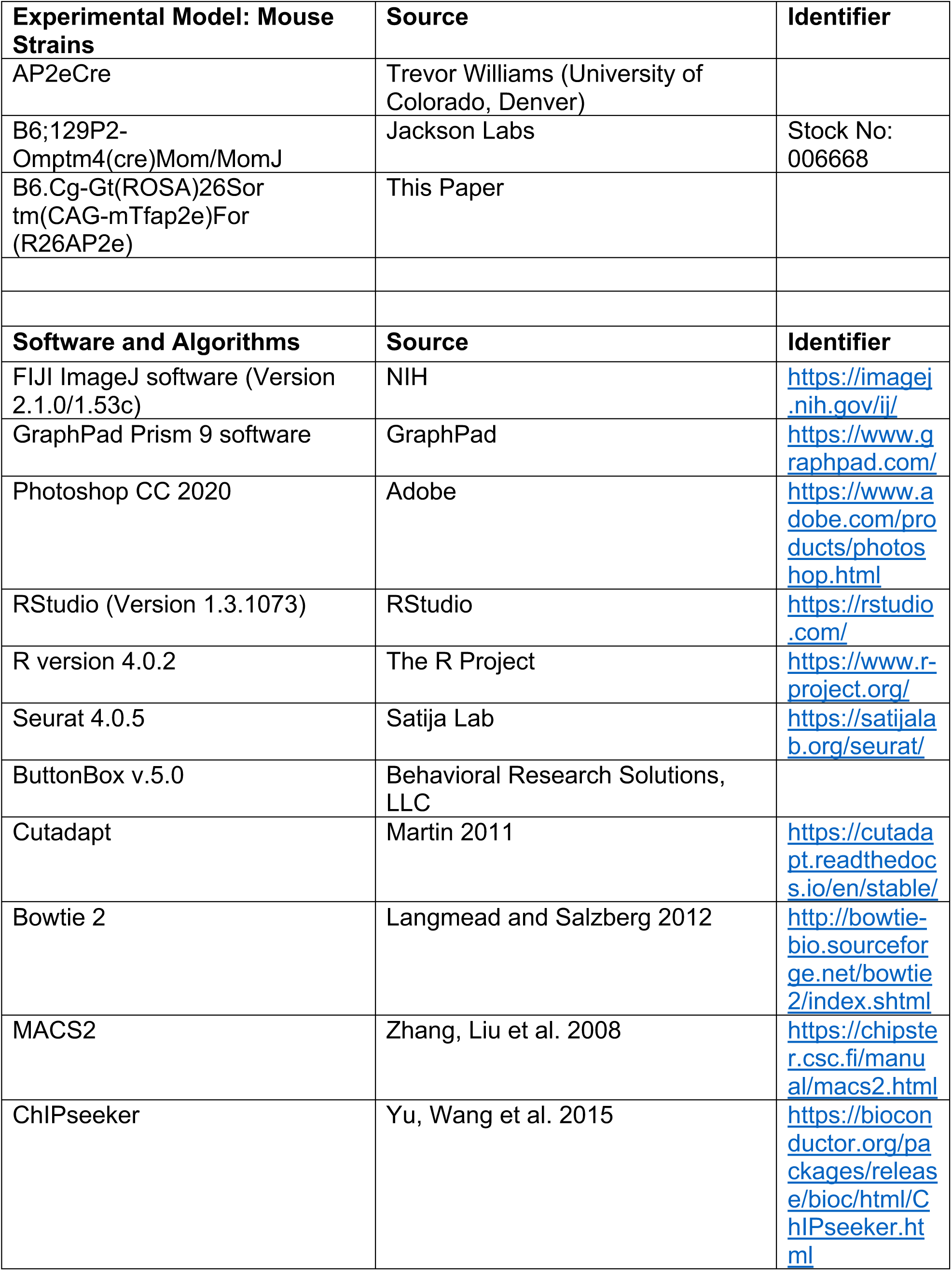
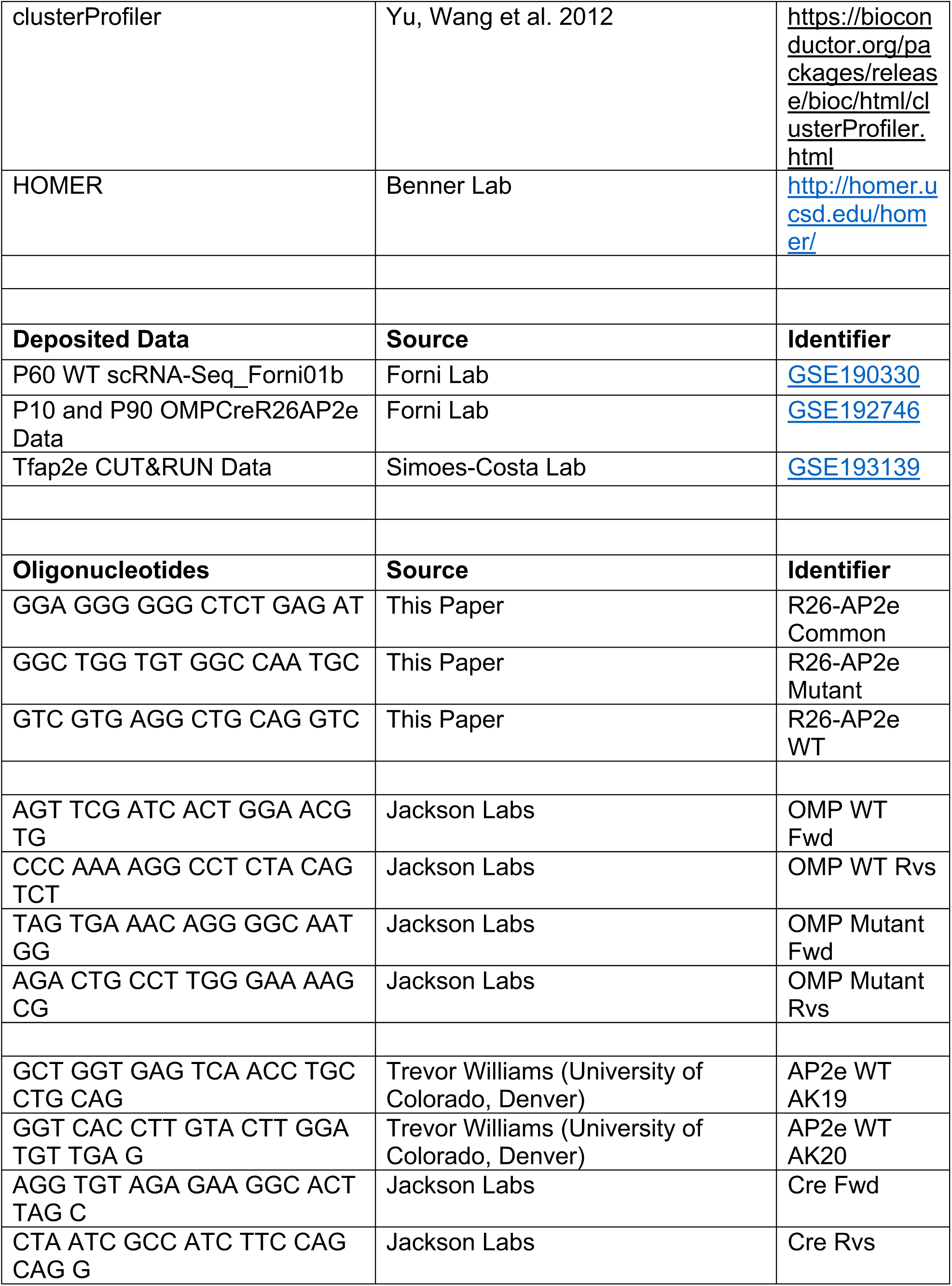
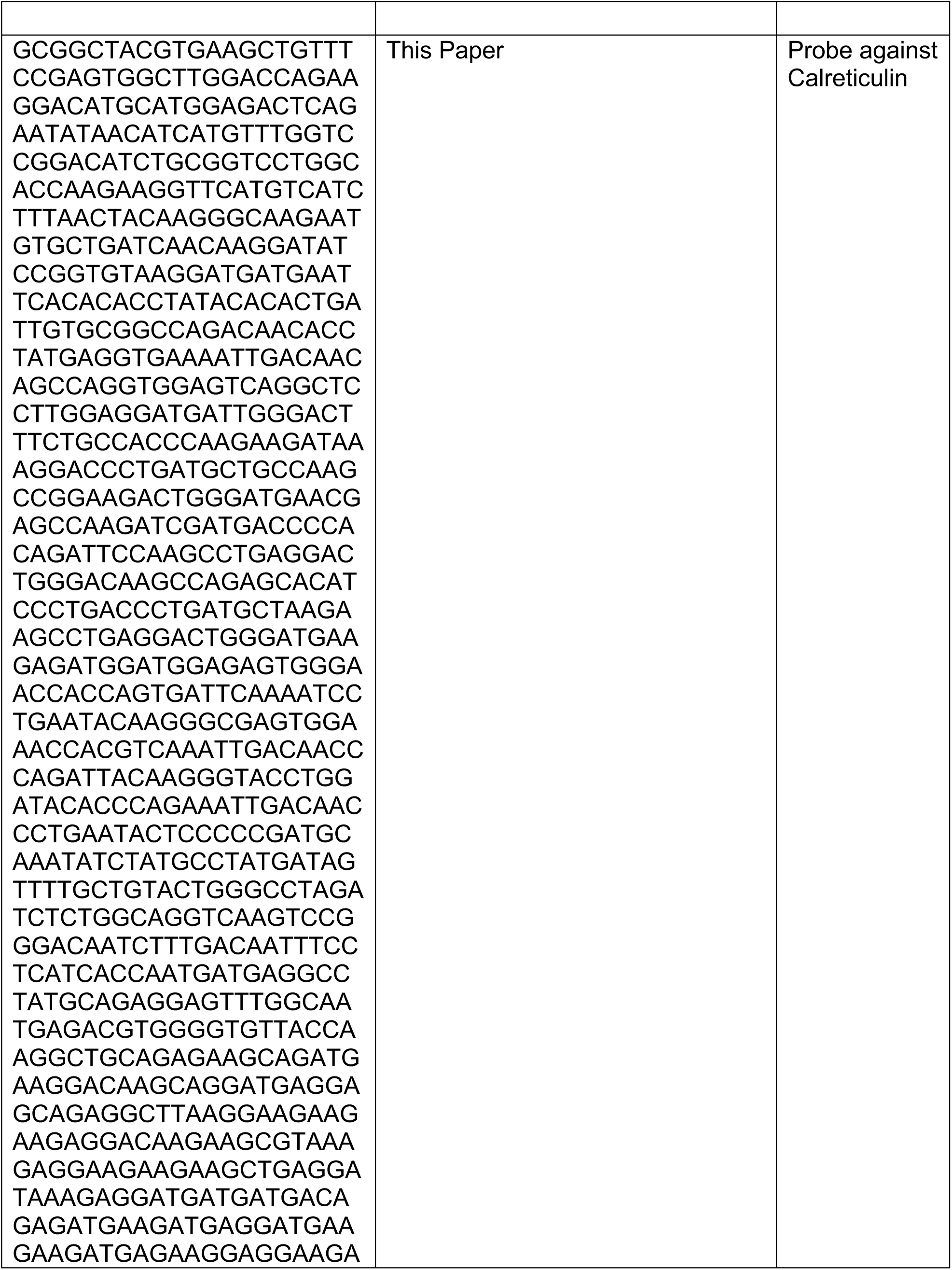
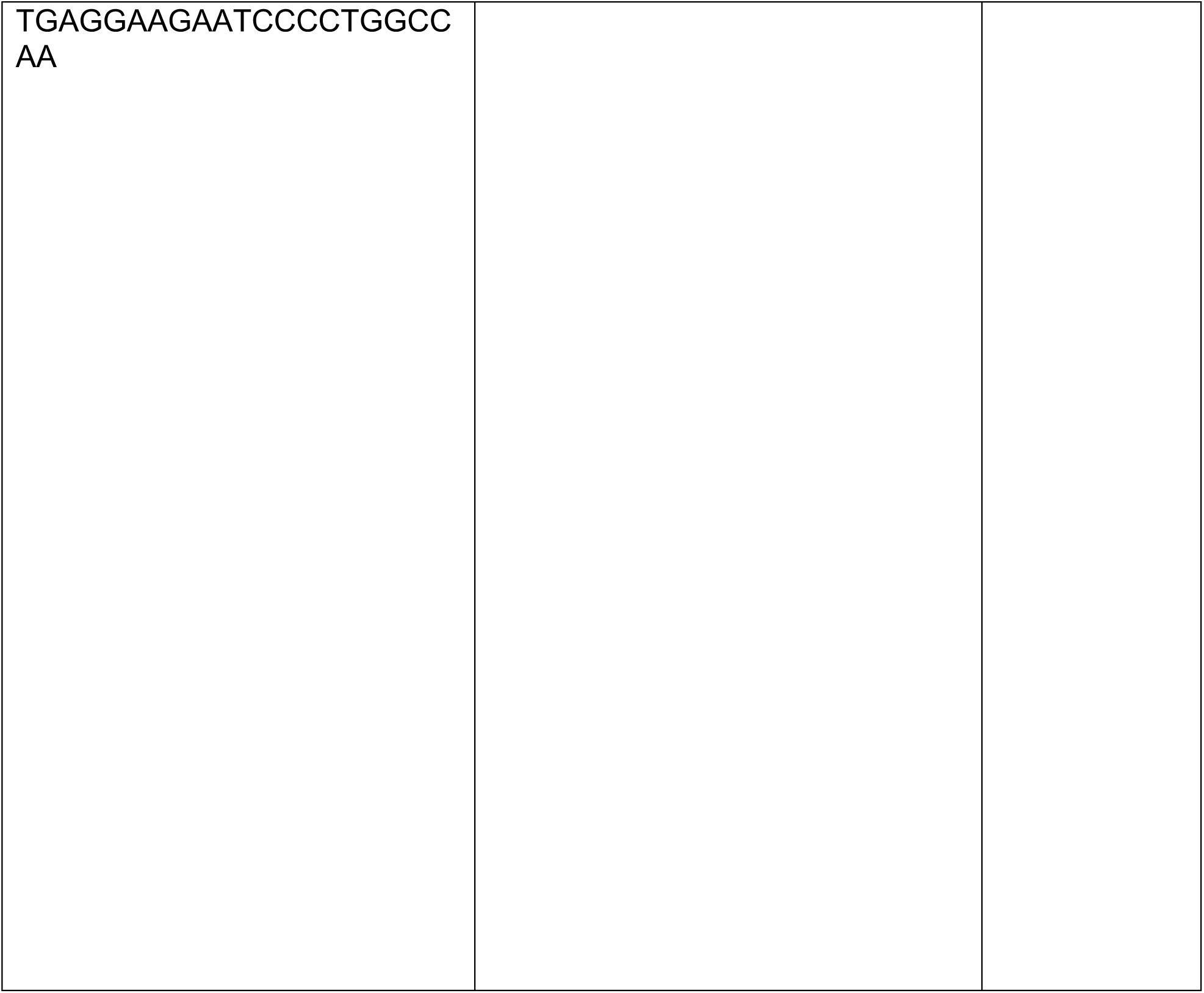
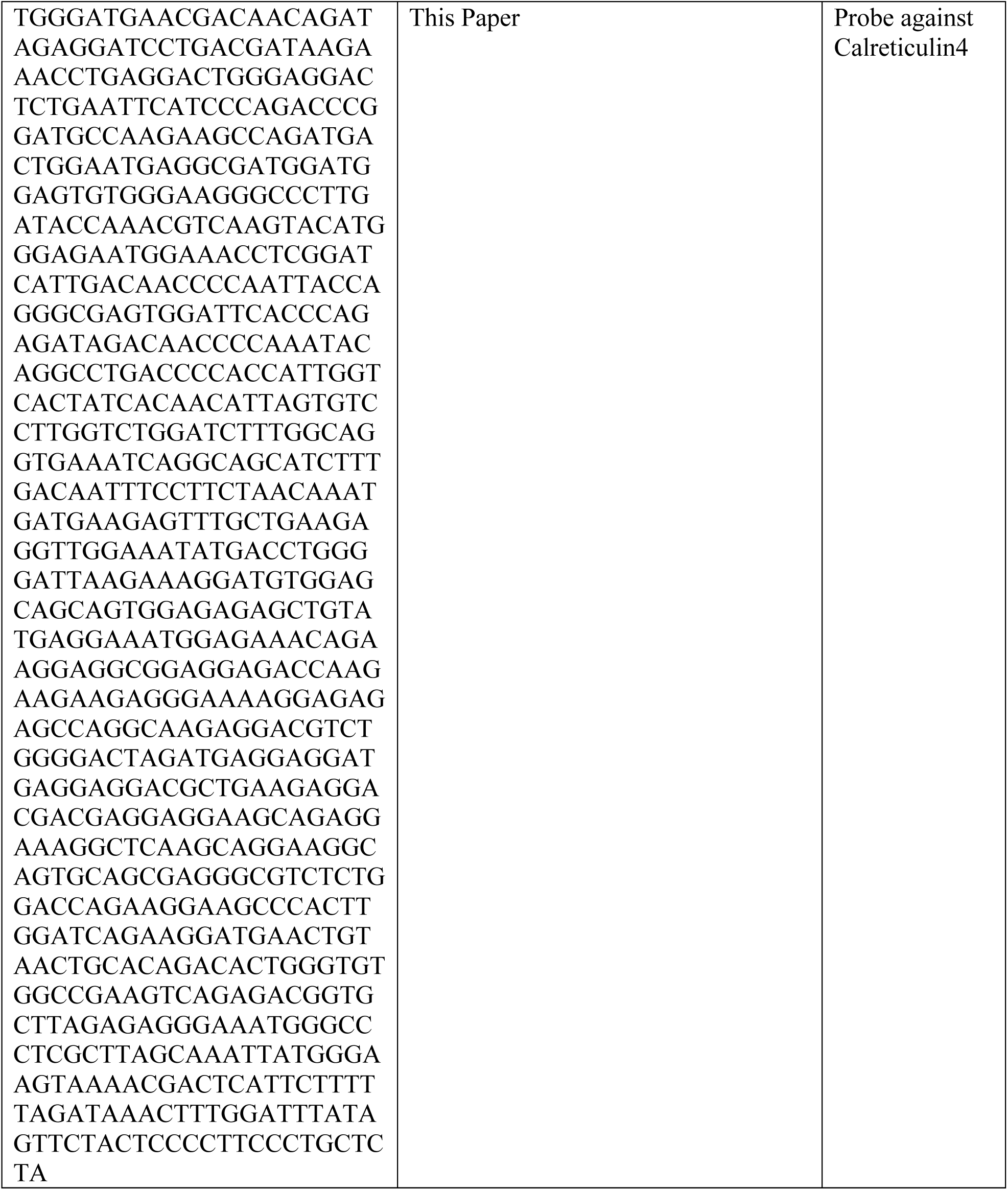

## RESOURCE AVAILABILITY

### Lead contact

Further information and requests for resources and reagents should be directed to and will be fulfilled by the lead contact, Paolo E. Forni.

### Materials availability

Mouse lines generated in this study will be deposited to Jackson Labs by the time of publication.

There are restrictions in availability of the antibody Rabbit anti-V2R2 which was obtained from the lab of Dr. Roberto Tornielli (University of Parma, Italy) and is not commercially available.

### Data and code availability

The Single-cell RNA-sequencing and CUT& RUN sequencing data discussed in this publication have been deposited at NCBI’s Gene Expression Omnibus and are publicly available as of the date of publication. Accession numbers are listed in the key resources table. This paper reports no original code. Any additional information required to reanalyze the data reported in this paper is available from the lead contact upon request.

## EXPERIMENTAL MODEL AND SUBJECT DETAILS

### Animals

The R26AP2ε mice were produced by Cyagen (Santa Clara, CA). on a C57B/6 background. The AP-2εCre line (*Tfap2e^tm1(cre)Will^*) was obtained from Dr. Trevor Williams, Department of Craniofacial Biology, University of Colorado. The R26AP-2ε (B6.Cg-Gt(ROSA)26Sor^tm(CAG-mTfap2e)For^) mouse line was produced through Cyagen on a C56BL/6 background. The OMPCre line (B6;129P2-*Omp^tm4(cre)Mom^*/MomJ) was obtained from Dr. Paul Feinstein (Hunter College, City University of New York) on a 129P2/OlaHsd background and backcrossed to a C57BL/6 background for 6 generations at the time of this study. The characterization and comparison of the rescue of the AP-2ε phenotype (AP-2εCreR26AP-2ε), AP-2ε KO, and wild-types were performed on a C57BL/6 background. OMPCre^+/-^/R26AP2ε^+/-^ mutant mice are viable. Genotyping of mutants was performed by PCR. Primers used are detailed in the Key Resources Table.

Mice were housed under a 12hr day/night cycle. Animals were collected/analyzed at P10, P21, and Adult (P60-P90) ages. For all morphological analyses both males and females were included unless otherwise specified. All mouse studies were approved by the University at Albany Institutional Animal Care and Use Committee (IACUC). Mouse lines generated in this study will be deposited to Jackson Labs by the time of publication.

## METHOD DETAILS

### Generation of the AP-2ε conditional Knock-In Model

The AP-2ε conditional Knock-In allele was generated by targeting the ROSA26 gene in C57BL/6 ES cells. The “CAG-loxP-stop-loxP-mouse Tfap2e CDS-polyA” cassette was cloned into intron 1 of ROSA26 in the reverse orientation. In the targeting vector, the positive selection marker (Neo) was flanked by SDA (self-deletion anchor) site, and DTA was used for negative selection. Mouse genomic fragments containing homology arms (Has) were amplified from BAC clone by using high fidelity Taq DNA polymerase and were sequentially assembled into a targeting vector together with recombination sites and selection markers.

The ROSA26 targeting construct was linearized by restriction digestion with AscI followed by phenol/chloroform extraction and ethanol precipitation. The linearized vector was transfected into C57BL/6 ES cells according to Taconic-Cyagen’s standard electroporation procedures and G418 resistant clones were selected for 24 hours post-electroporation. These were then screened for homologous recombination by PCR and characterized by Southern Blot analysis. Two separate clones, A2 and H_2_, were successfully transmitted to germline and characterized.

Genotyping for the R26AP-2ε mouse line was performed by PCR using R26-AP-2e Common (5’ GGAGGGGGGCTCTGAGAT 3’), R26-AP-2ε Mutant (5’ GGCTGGTGTGGCCAATGC 3’), R26-AP-2ε WT (5’ GTCGTGAGGCTGCAGGTC 3’) with expected bands at 552bp (Mutant) and 400bp (WT).

**Both OMPCre^+/-^ and wildtype mice are used as controls depending on availability during performed experiments.**

### Resident Intruder Test

The resident intruder assay was used to evaluate aggression in male mice of mutants and controls. Test subjects were housed with intact females for at least one week prior to testing. On the day of testing, all subjects (residents and intruders) were acclimated to the experimental environment for at least 30 minutes prior to the assay. Females were removed immediately before testing. Castrated C57B mice were swabbed with male whole urine immediately before being introduced into the resident male’s home cage. Interactions between isolated residents and intruders were recorded for 10min and videos were evaluated using ButtonBox v.5.0 (Behavioral Research Solutions, Madison WI, USA) software for the number and duration of attacks.

### Innate olfactory preference test

Adult mice were isolated for at least one week prior to testing. Individual mice were habituated to the experimental environment for at least 30 minutes, then to the test cage for an additional 2 minutes. After the habituation period, cotton swabs scented with either male or female whole urine was placed on either side of the test cage. The time spent sniffing each odorant was normalized to total investigation time.

### Neuronal activation in response to sex-specific odorants

Adult mice were isolated for at least one week prior to exposure to either soiled bedding from male or female mice for ∼90min then perfused with PBS and 3.7% formaldehyde in PBS, then collected to evaluate neuronal activation with immunohistochemistry against pS6.

### Tissue Preparation

Tissue collected at ages ≥P10 were perfused with PBS then 3.7% formaldehyde in PBS. Brain tissue was isolated at the time of perfusion and then immersion-fixed for 3-4 hours at 4°C. Noses were immersion fixed in 3.7% formaldehyde in PBS at 4°C overnight and then decalcified in 500mM EDTA for 3-4 days. All samples were cryoprotected in 30% sucrose in PBS overnight at 4°C, followed by embedding in Tissue-Tek O.C.T. Compound (Sakura Finetek USA, Inc., Torrance CA) using dry ice, and stored at -80°C. Tissue was cryosectioned using a CM3050S Leica cryostat at 16μm for VNOs and 20μm for brain tissue and collected on VWR Superfrost Plus Micro Slides (Radnor, PA) for immunostaining and in situ hybridization (ISH). All slides were stored at -80°C until ready for staining.

### Immunohistochemistry

For immunohistochemistry and immunofluorescence antigen retrieval was performed on slides that were submerged in citrate buffer (pH 6.0) above 95°C for at least 15min before cooling to room temperature, then permeabilized with and blocked in horse serum based blocking solution before transferring into primary antibodies overnight at 4°C. For immunohistochemistry slides were additionally incubated in an H_2_O_2_ solution (35mL PBS+ 15mL 100% Methanol + 500μL 30% H_2_O_2_) after antigen retrieval.

For chromogen-based reactions, staining was visualized with the Vectastain ABC Kit (Vector, Burlingame, CA) using diaminobenzidine (DAB) (Forni et al., 2011); sections were counterstained with methyl green and mounted with Sub-X mounting medium. For immunofluorescence species-appropriate secondary antibodies conjugated with either Alexa Fluor 488, Alexa Fluor 594, Alexa Fluor 568, Alexa Fluor 680 were used for immunofluorescence detection (Molecular Probes and Jackson ImmunoResearch Laboratories, Inc., Westgrove, PA). Sections were counterstained with 4’,6’-diamidino-2-phenylindole (DAPI) (1:3000; Sigma-Aldrich), and coverslips were mounted with FluoroGel (Electron Microscopy Services, Hatfield, PA).

Confocal microscopy pictures were taken on a Zeiss LSM 710 microscope. Epifluorescence pictures were taken on a Leica DM4000 B LED fluorescence microscope equipped with a Leica DFC310 FX camera. Images were further analyzed using FIJI/ImageJ software. Antibodies and concentrations used in this study are detailed in the Key Resources Table.

### In Situ Hybridization and RNAscope

Digoxigenin-labeled RNA probes were prepared by *in vitro* transcription (DIG RNA labeling kit; Roche Diagnostics, Basel, Switzerland). In situ hybridizations were performed on 16μm cryosections that were rehydrated in 1x PBS for 5min, fixed in 4% PFA in 0.1M phosphate buffer for 20min at 4°C, treated with 10μg/mL proteinase K (Roche) for 12min at 37°C, and then refixed in 4% PFA at 4°C for 20min. To inactivate the internal alkaline phosphatase, the tissue was treated with 0.2M HCl for 30min. Nonspecific binding of the probe to slides was reduced by dipping slides in 0.1M triethanolamine (pH 8.0)/0.25% acetic anhydride solution, then washed with 2x Saline-Sodium Citrate (SSC) buffer before incubating in hybridization solution for 2hrs at room temperature. Slides were then hybridized with 200μl of probe in hybridization solution at 65°C overnight in a moisture chamber. After hybridization, the slides were washed in 2x SSC, briefly, then in 1x SSC/50% formamide for 40min at 65°C. RNase A treatment (10μg/mL) was carried out at 37°C for 30min. The slides were then washed with 2x SSC then 0.2x SSC for 15min each at 65°C. Hybridization was visualized by immunostaining with an alkaline phosphatase conjugated anti-DIG (1:1000), and NBT/BCIP developer solution (Roche Diagnostics). After color reaction, the slides were put into 10mM Tris-HCl pH 8.0/1mM EDTA, rinsed in PBS and air dried before mounting with Sub-X mounting medium.

Single-molecule fluorescence in situ hybridization was performed using the RNAscope Multiplex Fluorescence v2 assay and probes (RNAscope Probe-Mm-Abca7-C4 #489021, RNAscope Probe-Mm-Gnai2 #868051, RNAscope Probe-Mm-Gnao1-E4-E6-C2 #444991, Rnascope Probe-Mm-Mt3-C3 #504061, RNAscope Probe-Mm-Meis2-C3 #436371) from ACDbio. The assay was performed on 16μm fixed-frozen P10-P11 mouse cryosections, following the manufacturer’s protocol.

### Single-Cell RNA Sequencing

The vomeronasal organs of OMPCre^+/-^ at P10 and OMPCre^+/-^/R26AP2ε^+/-^ at P10 and 3mo were isolated and dissociated into single-cell suspension using neural isolation enzyme/papain (NIE/Papain in Neurobasal Medium with 0.5mg/mL Collagenase A, 1.5mM L-cysteine and 100U/mL DNAse I) incubated at 37°C. The dissociated cells were then washed with HBSS and reconstituted in cell freezing medium (90% FBS, 10% DMSO). Cells were frozen from room temperature to -80°C at a -1°C/min freeze rate.

Single cell suspension was sent to SingulOmics for high-throughput single-cell gene expression profiling using the 10x Genomics Chromium Platform. Data were analyzed along with using Seurat 4.0.5. The scRNA-seq data discussed in this publication have been deposited in NCBI’s Gene Expression Omnibus and are accessible through GEO series accession number GSE192746 (https://www.ncbi.nlm.nih.gov/geo/query/acc.cgi?acc=GSE192746). We also utilized previously published data from (Katreddi et al., 2022), available through GEO series accession number GSE190330 (https://www.ncbi.nlm.nih.gov/geo/query/acc.cgi?acc=GSE190330).

## CUT&RUN

Cells frozen in 90% FBS/10%DMSO were thawed at 37°C and resuspended in CUT&RUN wash buffer (20mM HEPES pH7.5, 150mM NaCl, 0.5mM spermidine, plus Roche Complete Protease inhibitor, EDTA-free). CUT&RUN experiments were performed as previously described (Meers et al., 2019) with minor modifications. 0.025% digitonin was used for the Dig-wash buffer formulation. Antibody incubation was performed overnight at 4°C, followed by Protein A-MNase binding for 1 hour at 4°C. Prior to targeted digestion, cell-bead complexes were washed in low-salt rinse buffer (20mM HEPES pH7.5, 0.5mM spermidine, 0.025% digitonin, plus Roche Complete Protease inhibitor, EDTA-free) followed by targeted digestion in ice-cold high-calcium incubation buffer (3.5 mM HEPES pH 7.5, 10 mM CaCl_2_, 0.025% Digitonin) for 30 minutes at 0°C. Targeted digestion was halted by replacing the incubation buffer with EGTA-STOP buffer (170 mM NaCl, 20 mM EGTA, 0.025% digitonin, 20 µg/ml glycogen, 25 µg/ml RNase A), followed by chromatin release and DNA extraction. Protein AG–MNase was kindly provided by Dr. Steve Henikoff. A rabbit polyclonal Anti-TFAP2E antibody (Proteintech 25829-1-AP) was used at a concentration of 1:50 for CUT&RUN experiments.

## CUT&RUN library preparation

CUT&RUN libraries were prepared using the NEBNext ultra II DNA library prep kit (New England Biolabs E7645). Quality control of prepared libraries was conducted using an ABI 3730xl DNA analyzer for fragment analysis. Libraries were pooled to equimolar concentrations and sequenced with paired-end 37-bp reads on an Illumina NextSeq 500 instrument.

## QUANTIFICATION AND STATISTICAL ANALYSIS

### Quantification and statistical analyses of microscopy data

All data were collected from mice kept under similar housing conditions in transparent cages on a normal 12 hr. light/dark cycle. Tissue collected from either males or females in the same genotype/treatment group were analyzed together unless otherwise stated. Ages analyzed are indicated in text and figures. The data are presented as mean ± SEM. Prism 9.2.0 was used for statistical analyses, including calculation of mean values, and standard errors. Two-tailed, unpaired t-test were used for all statistical analyses, and calculated p-values <0.05 were considered statistically significant. Sample sizes and p-values are indicated as single points in each graph and/or in figure legends.

Measurements of VNE and cell counts were performed on confocal images or bright field images of coronal serial sections immunostained or in situ hybridizations for the indicated targets. In animals ≥P15, the most central 6-8 sections on the rostro-caudal axis of the VNO were quantified and averaged, and in animals ≥P0, the most medial 4-6 sections were quantified and averaged. Measurements and quantifications were performed using ImageJ 2.1.0 and Imaris. Statistical differences between genotypes were quantified with two-tailed unpaired t-test using Prism 9.2.0, (GraphPad Software, CA, USA). Microscopy data reported in this paper will be shared by the lead contact upon request.

### CUT&RUN data analysis

In processing CUT&RUN data, paired-end sequencing reads were trimmed using Cutadapt t(Martin, 2011) using the following arguments: “-a AGATCGGAAGAGCACACGTCTGAACTCCAGTCA -A AGATCGGAAGAGCGTCGTGTAGGGAAAGAGTGT --minimum-length=25”. Reads were aligned to the reference mouse mm10 assembly from the UCSC genome browser using Bowtie 2 (Langmead and Salzberg, 2012) using the following arguments: “--local --very-sensitive-local --no-unal --no-mixed --no-discordant -I 10 -X 1000”. BAM files were filtered with SAMtools to discard unmapped reads, those which were not the primary alignment, reads failing platform/vendor quality checks, and PCR/optical duplicates (-f 2 -F 780). Peak calling was performed using MACS2 (Zhang et al., 2008). Peak-gene annotation was done by mapping peaks to their closest annotated gene using the ChIPseeker R package(Yu et al., 2015). GO term analysis was performed in R using clusterProfiler (Yu et al., 2012). Motif enrichment analysis was performed using HOMER (Heinz et al., 2010). The data from this CUT&RUN experiment has been deposited into the NCBI’s Expression Omnibus and are accessible through GEO series accession number GSE193139 (https://www.ncbi.nlm.nih.gov/geo/query/acc.cgi?acc=GSE193139).

---

**Supplementary Figure S1:**
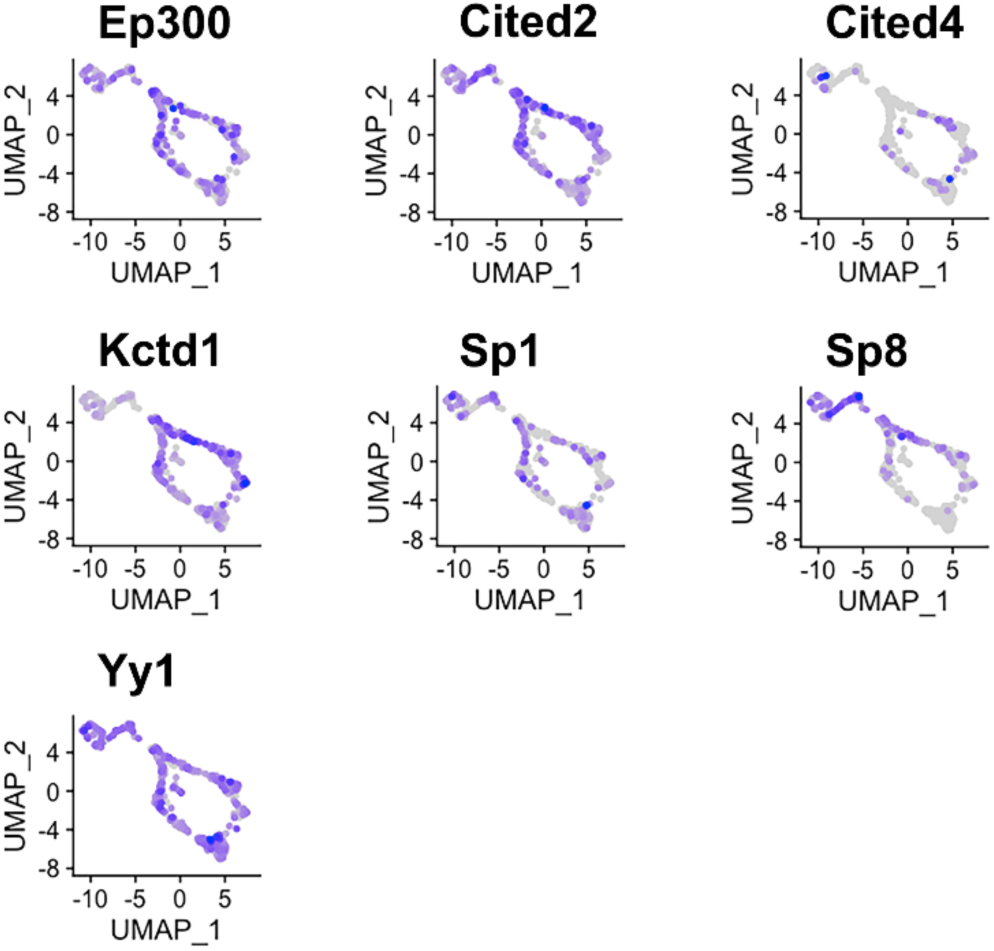
Expression of potential AP-2 co-factors in both apical and basal VSNs. Feature plots of known AP-2 family cofactors p300/CBP, Cited2, Cited4, Kctd1, Sp1, Sp8, Yy1 in P10 VSNs.

**Supplementary Figure S2:**
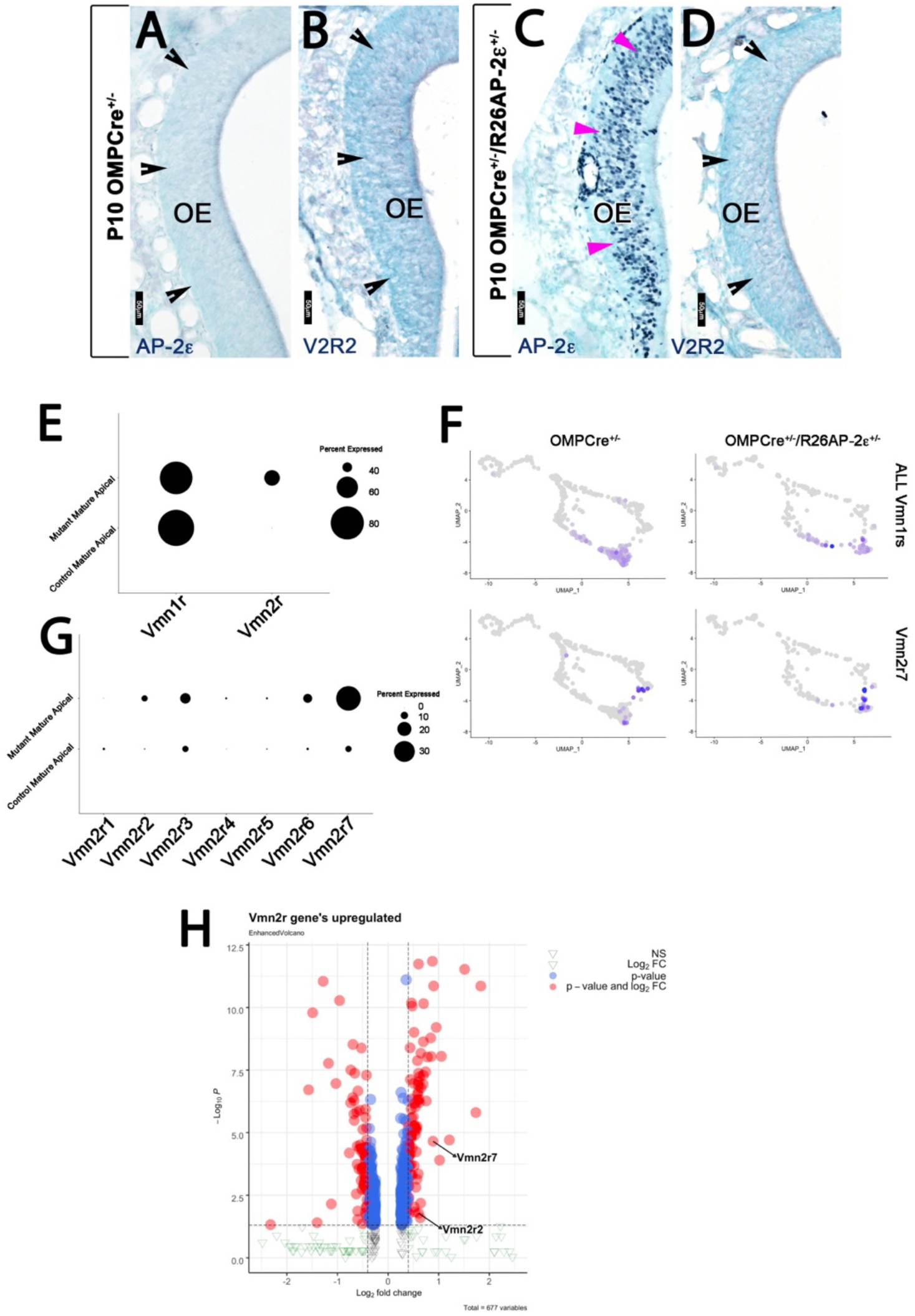
AP-2ε and V2R2 immunoreactivity in the MOE. P10 OMPCre^+/-^ controls(A,B) and OMPCre^+/-^/R26AP-2ε^+/-^ ectopic mutants (C,D). A,C) Immunohistochemistry against AP-2ε in the main olfactory epithelium (OE) shows no immunoreactivity in controls (black notched arrows) and ectopic AP-2ε expression in the mutants (magenta arrowheads). B,D) Immunohistochemistry against V2R2 in the main olfactory epithelium (OE) shows no expression (black notched arrowheads) in either control nor ectopic AP-2ε mutant.

**Supplementary Figure S3:**
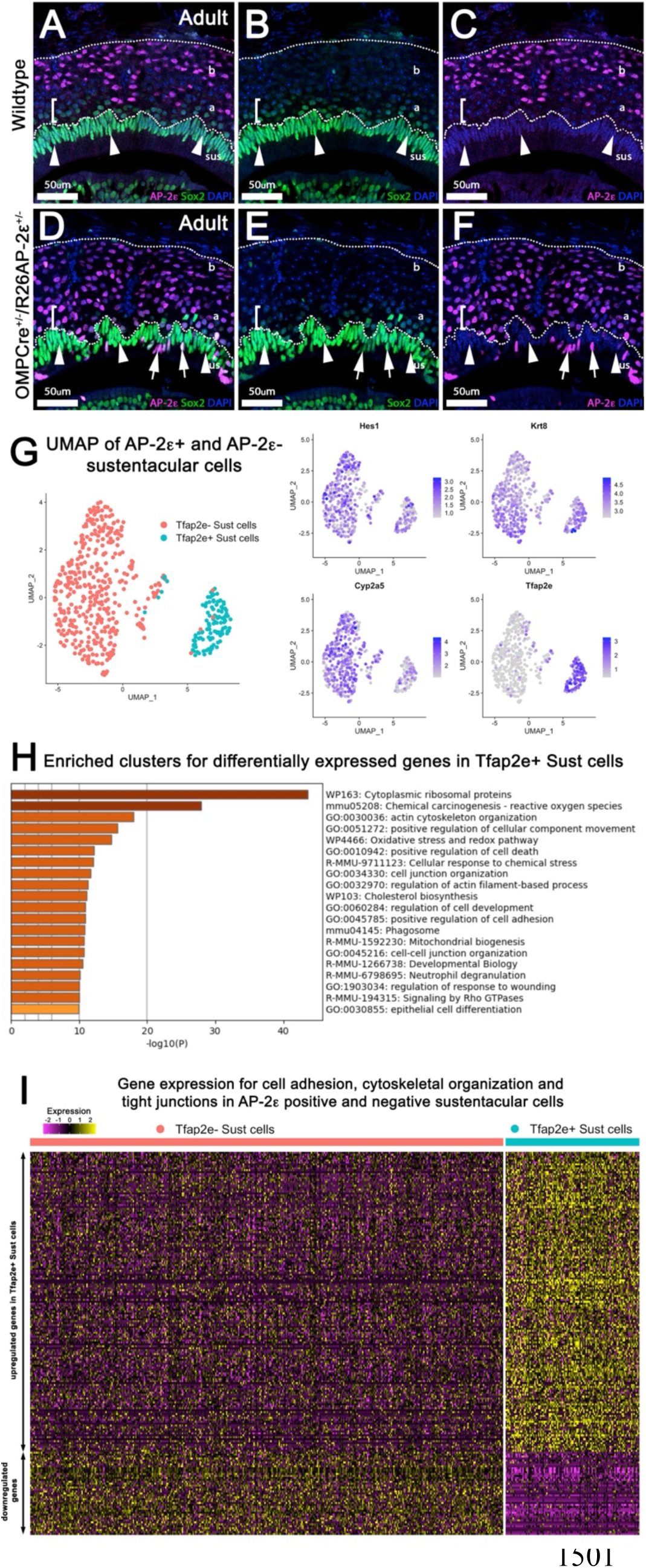
Analysis of Sustentacular Cells. A-F) Immunofluorescence on Wildtype (A-C) and OMPCre^+/-^/R26AP2ε^+/-^ mutants (D-F) against AP-2ε (magenta) and Sox2 (green) counterstained with DAPI (blue). A-C) A-C) WT controls and ectopic AP-2ε mutants (D-F) show sustentacular (sus) cells with high immunoreactivity for Sox2 and no AP-2ε expression. Apical (a) VSNs closest to the sustentacular cell layer show low immunoreactivity for Sox2 (brackets, arrowheads) with apical VSNs closer to basal (b) VSNs with no immunodetectable Sox2 (notched arrows). no colocalization between AP-2ε and Sox2+ sustentacular cells (sus). D-F) OMPCre^+/-^/R26AP2ε^+/-^ mice have some Sox2+ sustentacular cells that have ectopic AP-2ε expression (arrows). G) UMAPs of sustentacular cells from OMPCre^+/-^/R26AP2ε^+/-^ adults showed that AP2ε positive and negative clusters segregated from each other with accompanying feature plots showing the expression of sustentacular cell markers Hes1, Krt8, and Cyp2a5 along with ectopic Tfap2e/AP-2ε mRNA expression. H) Gene ontology analysis of Tfap2e+ sustentacular cells showed an enrichment of dysregulated genes related to cytoskeleton organization, cell-cell junction organization, cytoskeletal organization, and cytoplasmic ribosomal proteins. I) Heatmap of up- and down-regulated genes found related to cytoskeleton organization, cell-cell junction organization, and cytoskeleton organization in Tfap2e positive and negative sustentacular cells.

**Supplementary Figure S4.**
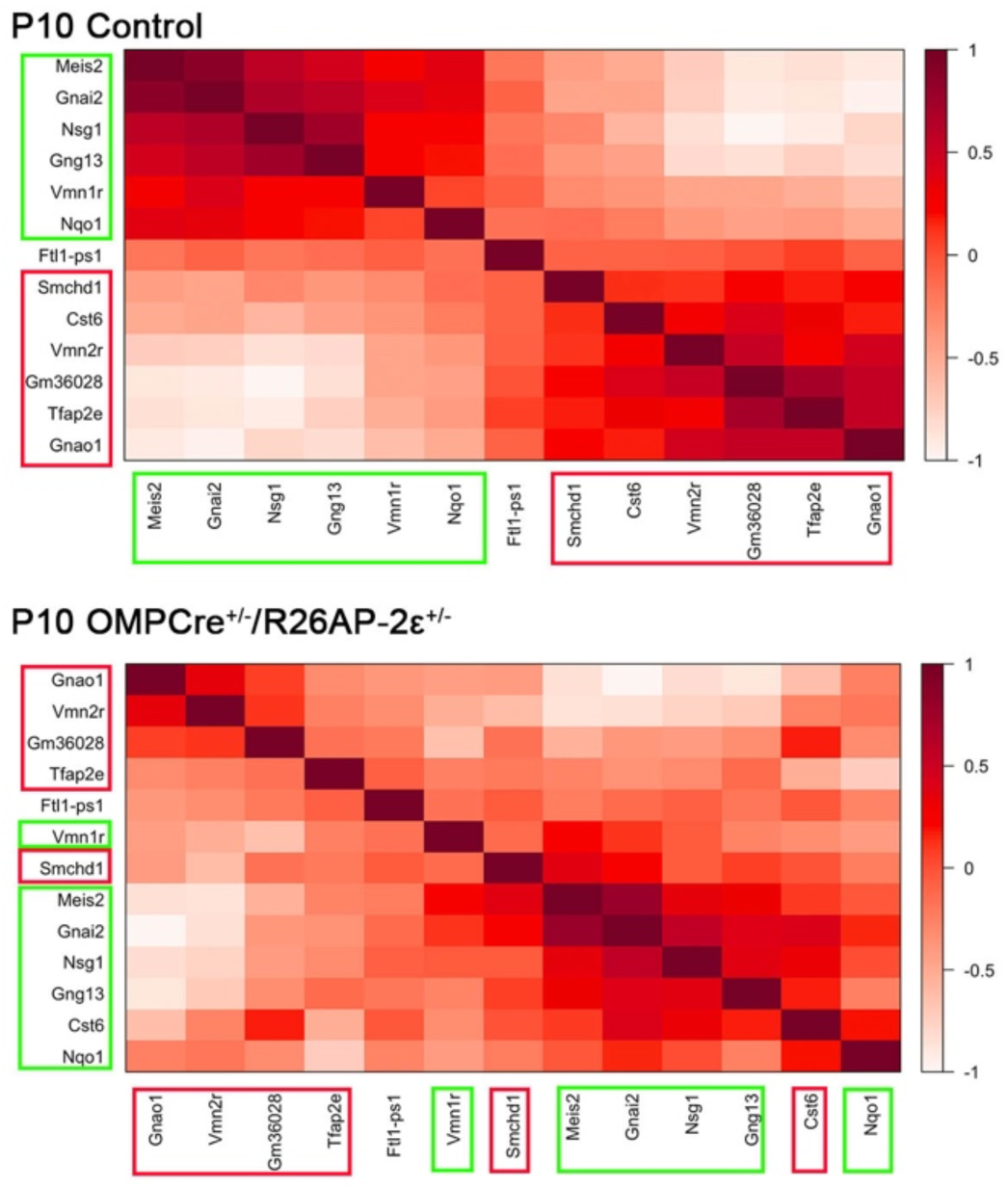
Gene Expression Correlation changes after ectopic AP- 2ε expression. In control animals, specific sets of genes are expressed with high correlation level in either apical VSNs (boxed in green) or basal VSNs (boxed in red). Genes with higher expression correlation are close to each other in either X and Y axis, the legend indicates higher correlation of expression with darker shade of red. In OMPCre^+/-^/R26AP2ε^+/-^ mutants, ectopic AP-2ε expression alters the cell type specific genes indicated by the different order of genes and different shades of red/expression correlation. These show that ectopic AP-2ε expression alters the normal correlation of apical and basal enriched genes.

**Supplementary Figure S5.**
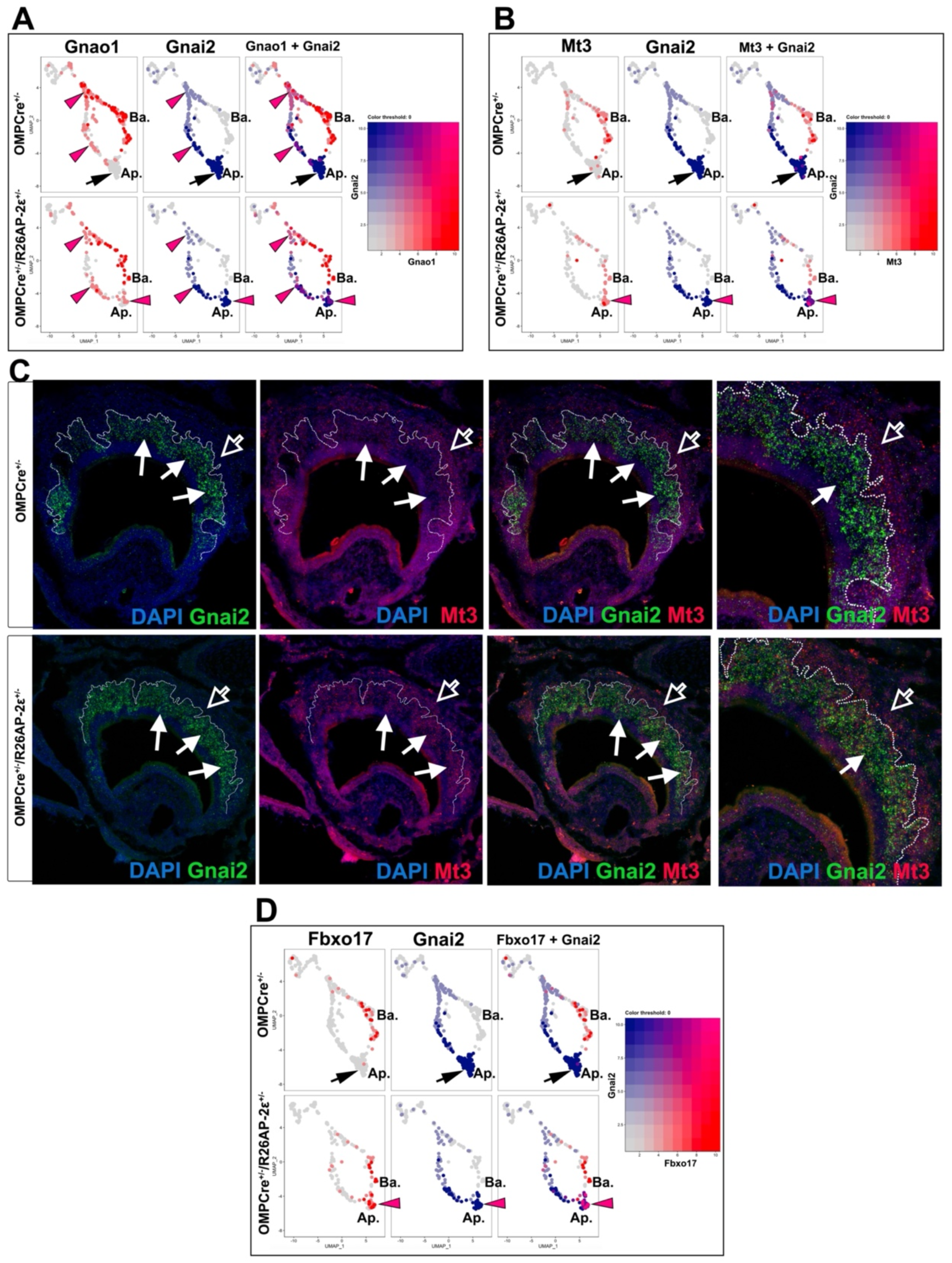
Ectopic expression in apical VSNs of indicated genes in OMPCre^+/-^/R26AP-2ε^+/-^ mutants. A) Feature plot showing ectopic expression of Gnao1 in OMPCre^+/-^/R26AP-2ε^+/-^ mutants. Black arrows indicate mature apical VSNs, Purple arrow heads indicate ectopic expression of Gnao1 in apical VSNs. B) Feature plot showing ectopic expression of Mt3 in OMPCre^+/-^/R26AP-2ε^+/-^ mutants. Black arrows indicate mature apical VSNs, Purple arrow heads indicate ectopic expression of Mt3 in apical VSNs. C) RNAscope analysis shows that Mt3 is mainly expressed in basal VSNs in OMPCre^+/-^ controls (arrow outline) but ectopically expressed in Gnai2+ apical VSNs in OMPCre^+/-^/R26AP-2ε^+/-^ mutants (arrows). D) Feature plot showing ectopic expression of Fbxo17 in OMPCre^+/-^/R26AP-2ε^+/-^ mutants. Black arrows indicate mature apical VSNs, Purple arrow heads indicate ectopic expression of Fbxo17 in apical VS

**Supplementary Figure S6:**
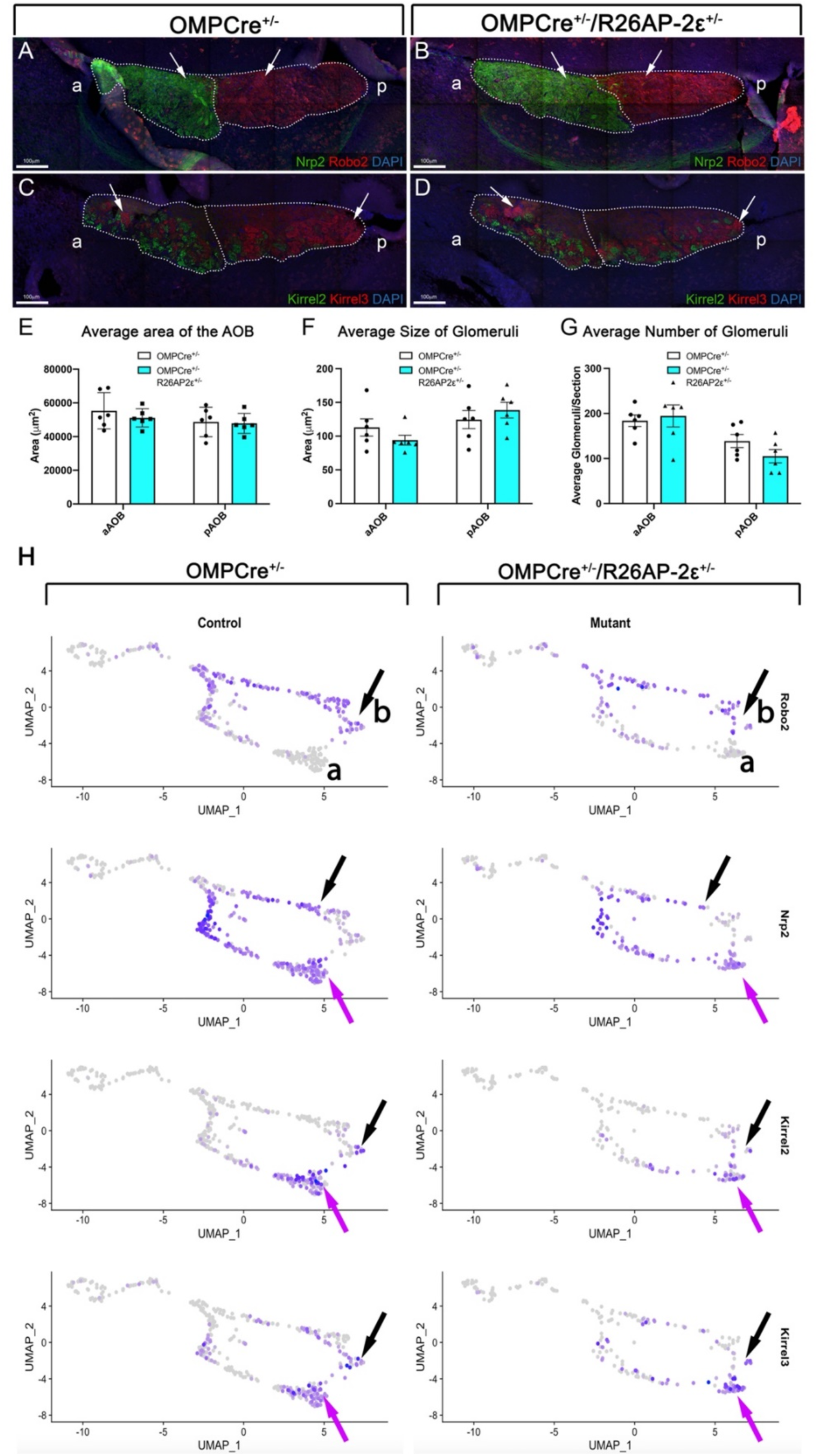
Analysis of the AOBs from adult WT and OMPCre^+/-^/R26AP2ε^+/-^ mice. A,B) Immunofluorescence against Nrp2 (green) and Robo2 (red) and counterstained with DAPI (blue) showed that the expression of guidance cue molecules in WT (A) and OMPCre^+/-^/R26AP2ε^+/-^ mutant (B) mice was comparable with distinct anterior and posterior AOB regions (arrows). C,D) Immunofluorescence against Kirrel2 (green) and Kirrel3 (red) and DAPI (blue) counterstain on WT (C) and OMPCre^+/-^/R26AP2ε^+/-^ mutant mice (D) AOBs showed that the glomerular organization and morphology was comparable between genotypes (arrows). E) Quantifications of the average area of the AOB was not significantly different across genotypes. Analysis of the glomeruli size (F) and number (G) using IF against Kirrel2 and Kirrel3 showed no significant changes across genotypes. N=6 for OMPCre^+/-^ controls and OMPCre^+/-^/R26AP2ε^+/-^ mutants (E-G).

**Supplementary Figure S7:**
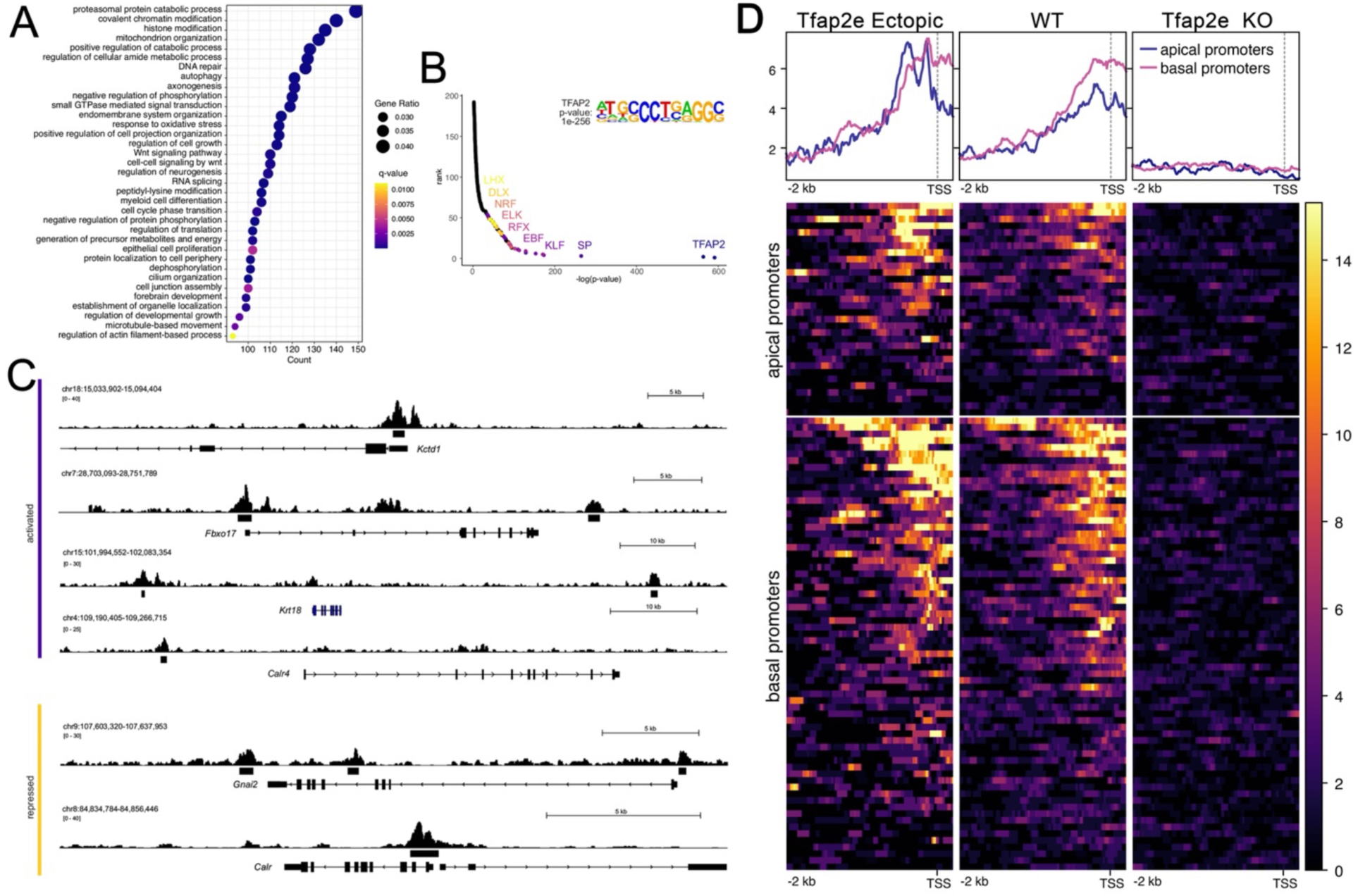
A) Dot plot of enriched GO terms show many putative AP-2ε target genes being chromatin or histone modifiers. B) Motif enrichment analysis validated our analysis as Tfap2 was the top motif found at putative binding sites. It also yielded TF family motifs of potential cofactors including SP, KLF, EBF, RFX, ELK, NRF, DLX, and LHX transcription factor families. C) Tracks from CUT&RUN sequencing where AP-2ε peaks are found in the promoter regions of activated (purple) and repressed (yellow) genes. D) Promotor plots of AP-2ε binding in ectopic mutants, WTs and KOs. In WTs, there is more Tfap2e binding at basal promotors compared to apical, however, in ectopic mutants, apical and basal binding is similar due to Tfap2e ectopic expression. Tfap2e KOs show little to no signal.

**Supplementary Figure S8:**
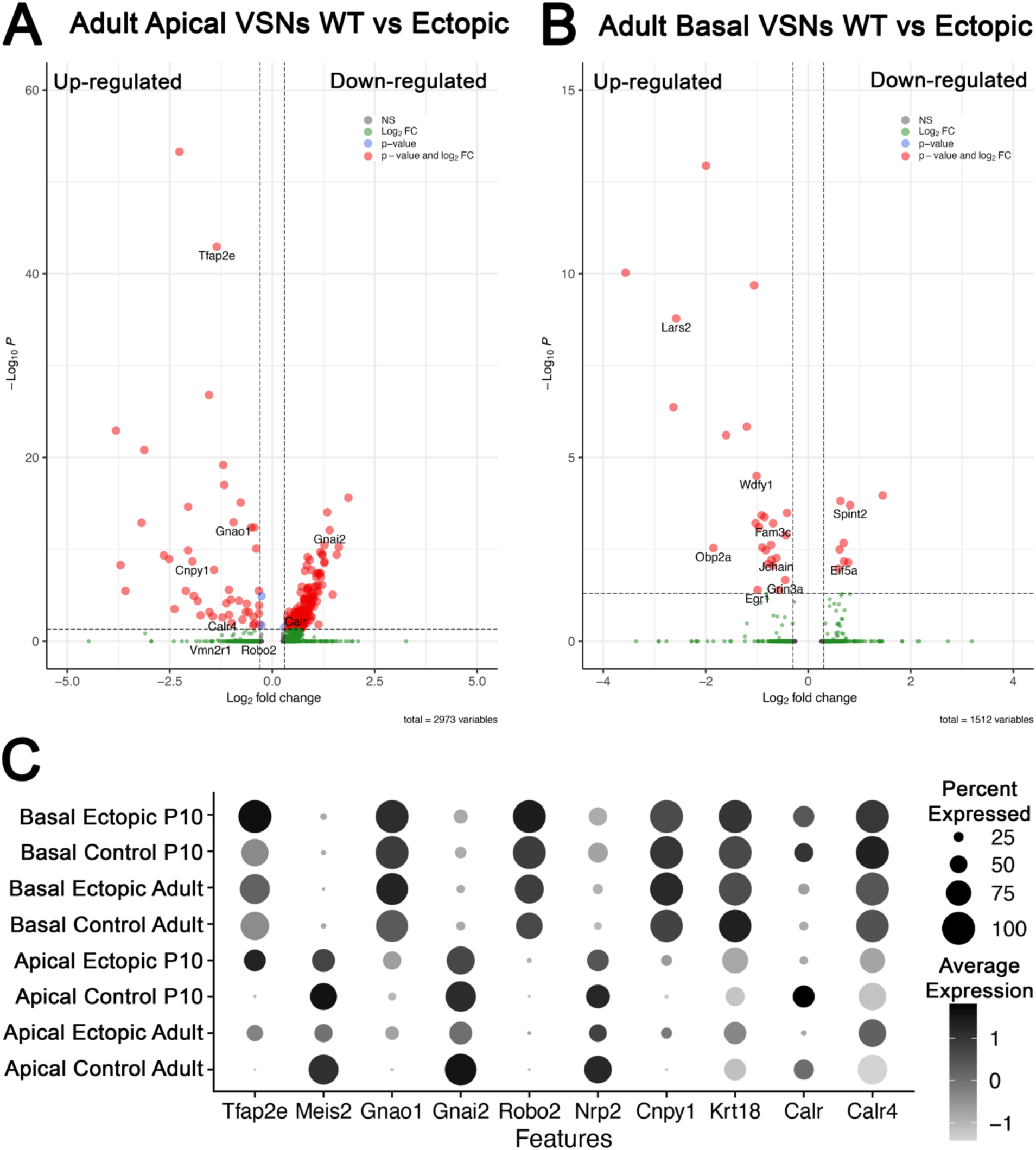
Differential gene expression analysis from scSeq in adult animals of apical and basal populations scSeq in Adults WT Controls and OMPCre^+/-^/R26AP2ε^+/-^ VNOs. A) Volcano Plot of significantly up- and down-regulated genes in apical VSNs in adult WT and OMPCre^+/-^/R26AP2ε^+/-^ mice show dysregulation of many key basal genes and down-regulation of apical-enriched genes. B) Volcano Plot of significantly up and downregulated genes in basal VSNs in adult WT and OMPCre^+/-^/R26AP2ε^+/-^ mice show that some genes may respond to AP-2ε regulation in a dose-dependent manner. C) Dot plot of canonical genes expressed by apical and basal VSNs and key genes identified in scSeq analysis for P10 and Adult Control and Mutant mice.

**Supplementary Table S1:**
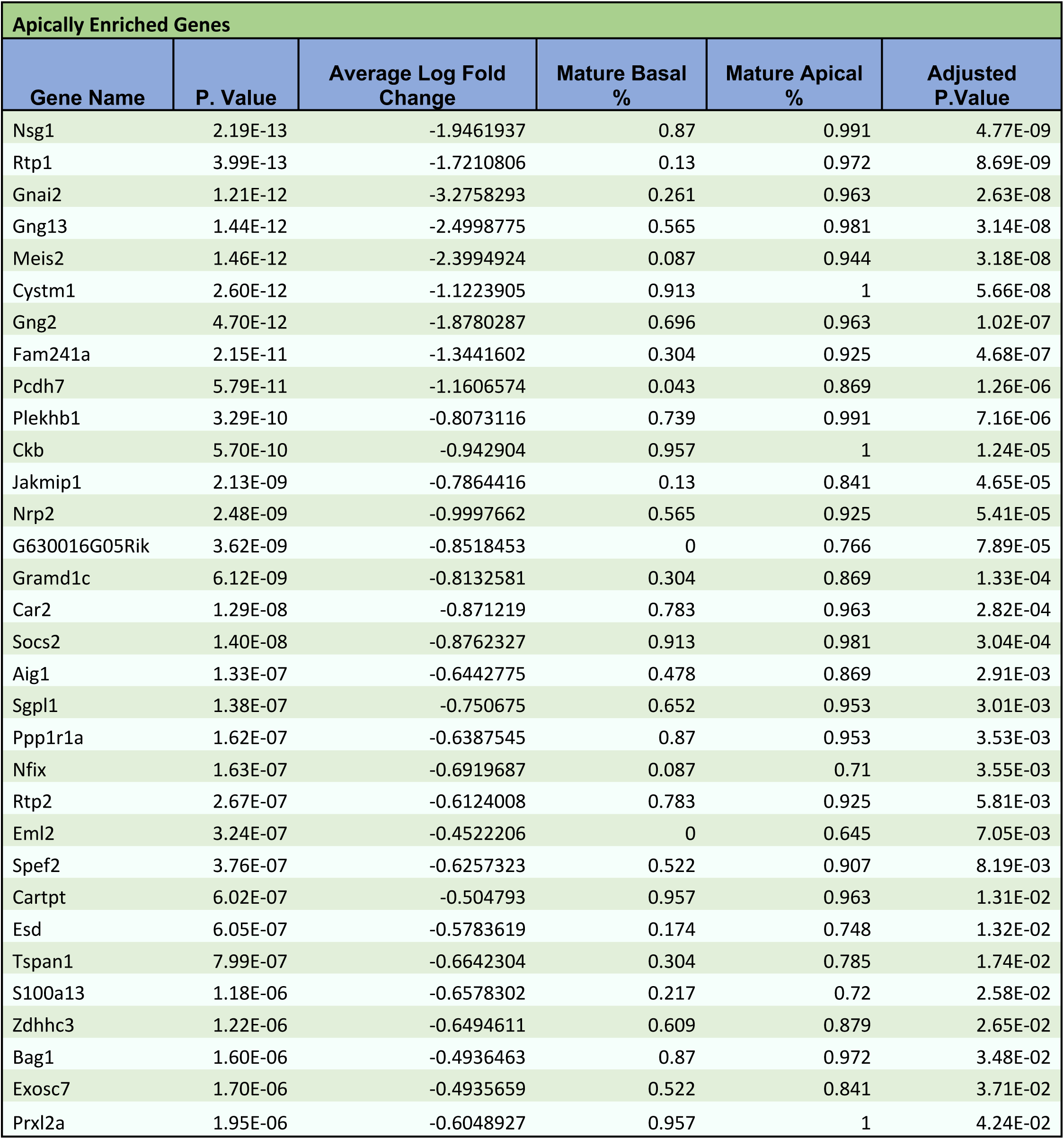
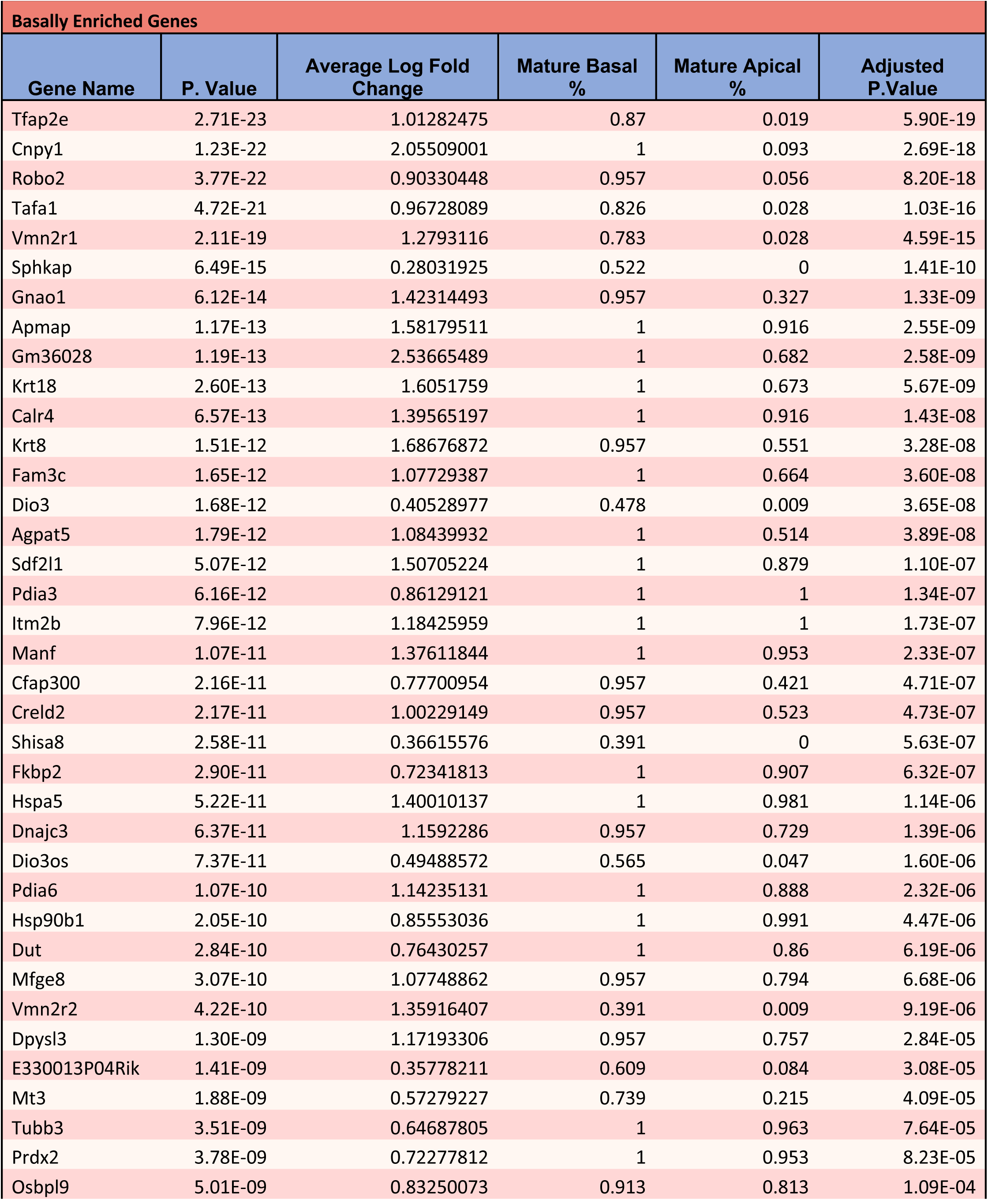
Differentially expressed genes in mature apical and basal VSNs.

**Supplementary Table S2:**
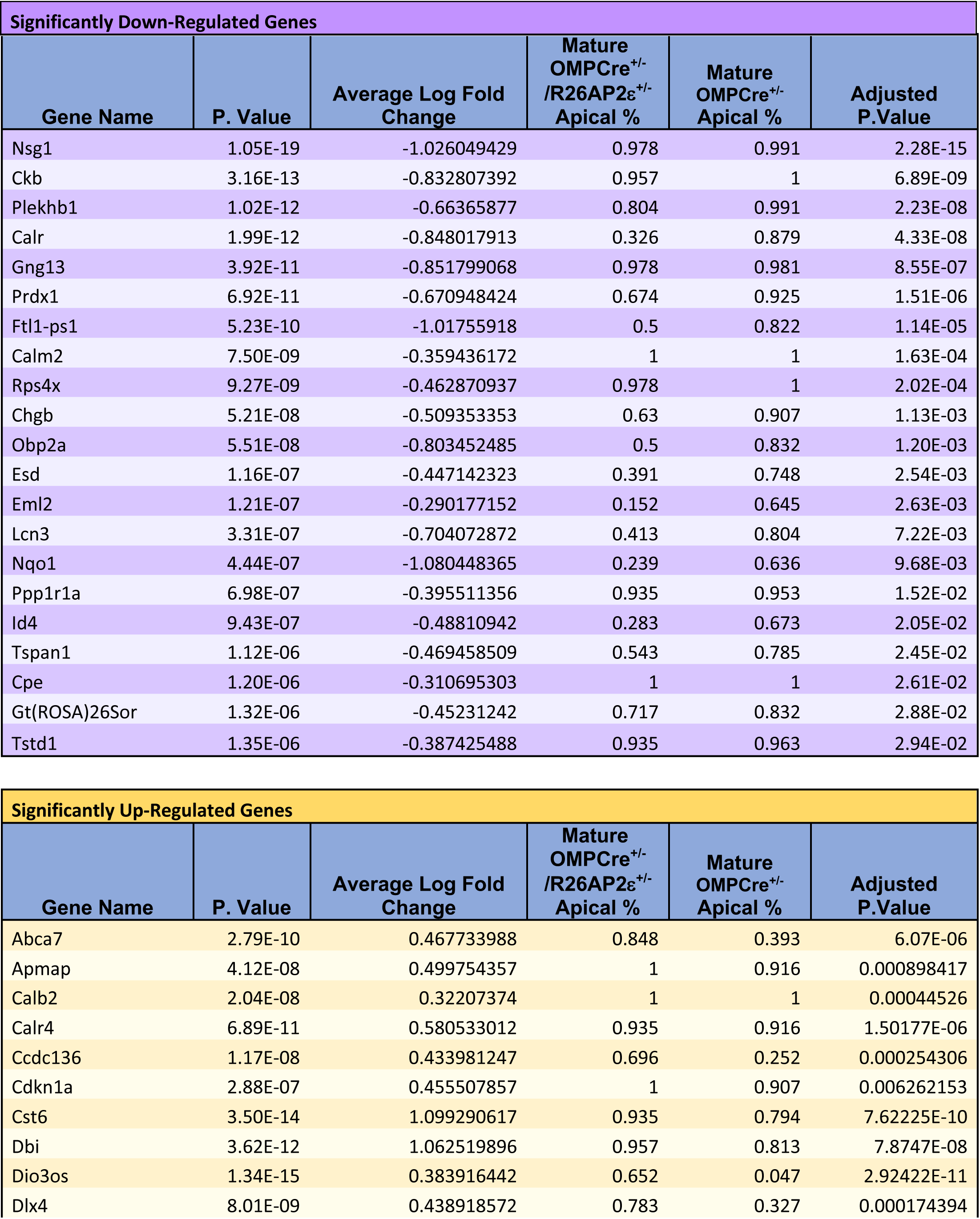
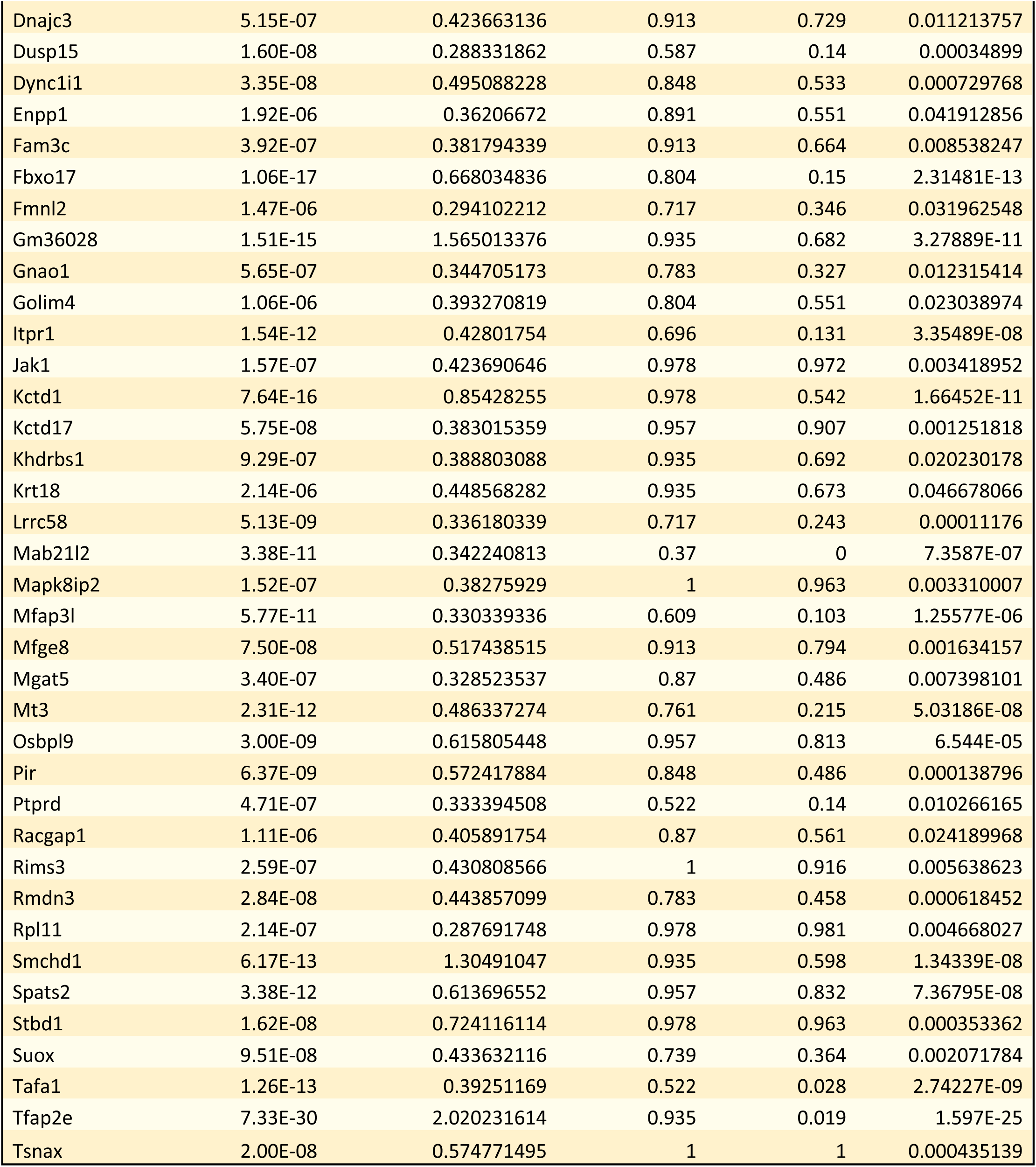
Significantly dysregulated genes when Tfap2e/AP-2ε is ectopically expressed in apical VSNs.

